# Breast cancer progression and metastasis to lymph nodes reveals cancer cell plasticity and MHC class II-mediated immune regulation

**DOI:** 10.1101/2022.10.30.514441

**Authors:** Pin-Ji Lei, Ethel R. Pereira, Patrik Andersson, Zohreh Amoozgar, Jan Willem Van Wijnbergen, Meghan O’Melia, Hengbo Zhou, Sampurna Chatterjee, William W. Ho, Jessica M. Posada, Ashwin Srinivasan Kumar, Satoru Morita, Charlie Chung, Ilgin Ergin, Dennis Jones, Peigen Huang, Semir Beyaz, Timothy P. Padera

**Affiliations:** Edwin L. Steele Laboratories, Department of Radiation Oncology, Massachusetts General Hospital Cancer Center, Massachusetts General Hospital and Harvard Medical School, Boston, MA 02114, USA; Cold Spring Harbor Laboratory, Cold Spring Harbor, New York 11724, USA; Department of Pathology and Laboratory Medicine, Boston University School of Medicine, 670 Albany Street, Boston, Massachusetts 02118, USA; Harvard–MIT Division of Health Sciences and Technology, Massachusetts Institute of Technology, Cambridge, MA, USA; Koch Institute for Integrative Cancer Research, Massachusetts Institute of Technology, Cambridge, MA 02142, USA

**Keywords:** breast cancer, lymph node metastasis, single-cell RNA sequencing, epithelial-to-mesenchymal transition, cancer cell heterogeneity, immune evasion

## Abstract

Tumor-draining lymph nodes are critical sites for generating tumor antigen-specific T cells and are associated with durable immune responses. However, lymph nodes are often the first site of metastasis and lymph node metastases portend worse outcomes. Through cross-species single cell gene expression analysis of breast cancer progression and metastasis to lymph nodes, we uncovered features that define the heterogeneity, plasticity, and immune evasion of cancer cells. Notably, a subpopulation of metastatic cancer cells in the lymph node were marked by high levels of MHC class II (MHC-II) gene expression both in mice and humans. Mechanistically, the IFN-γ and JAK/STAT signaling pathways mediate MHC-II expression in cancer cells. Ablation of IFNGR1/2 or CIITA, the transactivator of MHC-II, in cancer cells prevented tumor progression. Interestingly, MHC-II+ cancer cells lacked co-stimulatory molecule expression, engendered the expansion of regulatory T cells and blunted CD4+ effector T cells in the tumor-draining lymph nodes and favor tumor progression. Overall, our data suggests that cancer cell plasticity during breast cancer progression and metastasis to lymph nodes endows metastatic cells with the ability to avoid immune surveillance. These data provide the basis for new opportunities to therapeutically stimulate anti-cancer immune responses against local and systemic metastases.

## Introduction

In multiple solid tumors—including breast carcinomas, head and neck carcinomas, colon cancer and melanoma—the first sites of metastasis are often the tumor draining lymph nodes (TDLNs)^1–4^. The presence of lymph node metastasis is strongly correlated with poor prognosis and guides treatment strategies^5–9^. Recently, we and others independently showed that cancer cells in some lymph node metastases (LNMs) can escape the lymph nodes and disseminate to distant organs ^10,11^.

TDLNs are also the first opportunity for the immune system to experience tumor antigens and generate an anti-tumor immune response. Emerging studies have shown that TDLNs contain TCF1+ stem-like CD8 T cells and tumor-specific resident memory CD8 T cells that maintain active anti-cancer immune responses^12,13^. Further, tumor-specific PD-1+ T cells are enriched in TDLNs, PD-1/PD-L1 interactions occur frequently in TDLNs and blocking PD-L1 in TDLNs elicited effective anti-tumor immunity^14^. Paradoxically, metastatic cancer cells can survive and grow in TDLNs, which should have an immune response trained to attack these cancer cells. How do metastatic cancer cells grow and survive in lymph nodes remain understudied.

Recent studies have shown the metastatic and tumor draining lymph nodes are immunosuppressed by the cancer^15–18^. We have shown that there is a lack of lymphocyte infiltration into metastatic lesions in lymph nodes, which may limit immune activation^19^. In addition, the down regulation of MHC-I on some cancer cells and the presence of regulatory T cells (Tregs) provide additional evidence for an immune suppressed environment in metastatic lymph nodes^20^. Collectively, these findings underscore a critical role for TDLNs in anti-tumor immunity and the role metastatic lymph nodes play in cancer progression and immune suppression.

As metastatic cancer cells arrive in the lymph node, they experience and respond to a new microenvironment. Studies have shown that the lymph node microenvironment promotes metabolic remodeling of cancer cells by up-regulation of genes related to lipid and fatty acid metabolism, which could lead to the selection of more aggressive cancer cells in metastatic lymph nodes (metLNs) compared to the primary tumor^21,22^. Several studies have also shown that cancer cells disseminating from lymph nodes can acquire properties that allow them to better cope with stresses in the blood during subsequent metastasis^23^ and make themselves less susceptible to NK cell killing^20^. Despite the cancer cell heterogeneity that is generated by the process of metastasis, currently, most metastatic tumors are diagnosed and treated based on pathological characteristics of the primary tumor growing in its native microenvironment. This can lead to differential treatment responses of primary tumor and metastases that grow in lymph nodes^24^ or distant sites. A comprehensive understanding of the biological differences between primary tumors and LNMs that occur as a part of cancer progression is critical to optimize strategies for treatment of metastatic disease. Further, how cancer cells can grow and survive in lymph nodes by evading immune surveillance warrants further investigation^20^.

In this study, we measured the transcriptional profiles of cells in the primary tumor and metLNs at single-cell resolution in a murine model of spontaneous LNMs from orthotopic breast cancer. Not surprisingly, we found heterogeneity in the cancer cells of the primary tumor, including a majority of primary cancer cells that were *EpCAM, Vim*, and *Twist1* triple positive epithelial-to-mesenchymal transition (EMT)-hybrid cells. In metLNs, we found most cancer cells either displayed mesenchymal-like or epithelial-like phenotypes. Single-cell trajectory analysis suggested that mesenchymal-to-epithelial transition (MET) of cancer cells occurred in metLNs. The epithelial-like cancer cells showed elevation of MHC-II genes and lacked expression of immune co-stimulatory molecules. We also observed the presence of MHC-II on a population of cancer cells in human breast cancer LNMs. Further, we found that MHC-II expression on 4T1 cancer cells can suppress CD4+ effector T cell proliferation and enhance Treg proliferation *in vitro* and *in vivo*. Additionally, we also found that Treg cells in the metLNs displayed an enhanced immunosuppressive phenotype. Our study reveals a novel mechanism by which cancer cells adapt and induce an immunosuppressive environment in metLNs in order to evade attack by the immune system. These findings may provide strategies to treat LNMs and prevent their spread to distant organs.

## Results

### The single-cell atlas of murine breast cancer progression to lymph node metastasis

To understand the mechanism of how breast cancer cells invade and survive in the lymph nodes, we characterized cancer cells in both primary tumors and metLNs using 4T1 basal-like triple negative murine breast cancer that develops spontaneous lymph node metastasis from orthotopically implanted tumors in immunocompetent, syngeneic hosts^25^. We collected the primary tumor and metLN from the same mouse and performed single-cell RNA sequencing (n=3) (**Figure 1A**). In the primary tumor samples, we identified 7428 single cells that grouped into 22 unique cell clusters (**Figure S1A**). Next, we identified the most up-regulated genes in each cluster to annotate the individual cell types. Around 70% of the cells in the primary tumor microenvironment (TME) are immune cells (**Figure 1B and S1B**), as shown by the expression of *Ptprc*—the gene encoding CD45. The myeloid-derived monocytes and macrophages are the largest population of immune cells in the TME. In the metLNs, we identified 6029 single cells that grouped into 23 cell clusters. According to these marker genes, we found a rich diversity of cells in the metLNs, including cancer cells, conventional CD4 T cells, CD8 T cells, Treg cells, B cells, NK cells, macrophages, dendritic cells, plasmacytoid dendritic cells and lymph node stromal cells (**Figure 1B and S1C-D**).

**Figure1:**
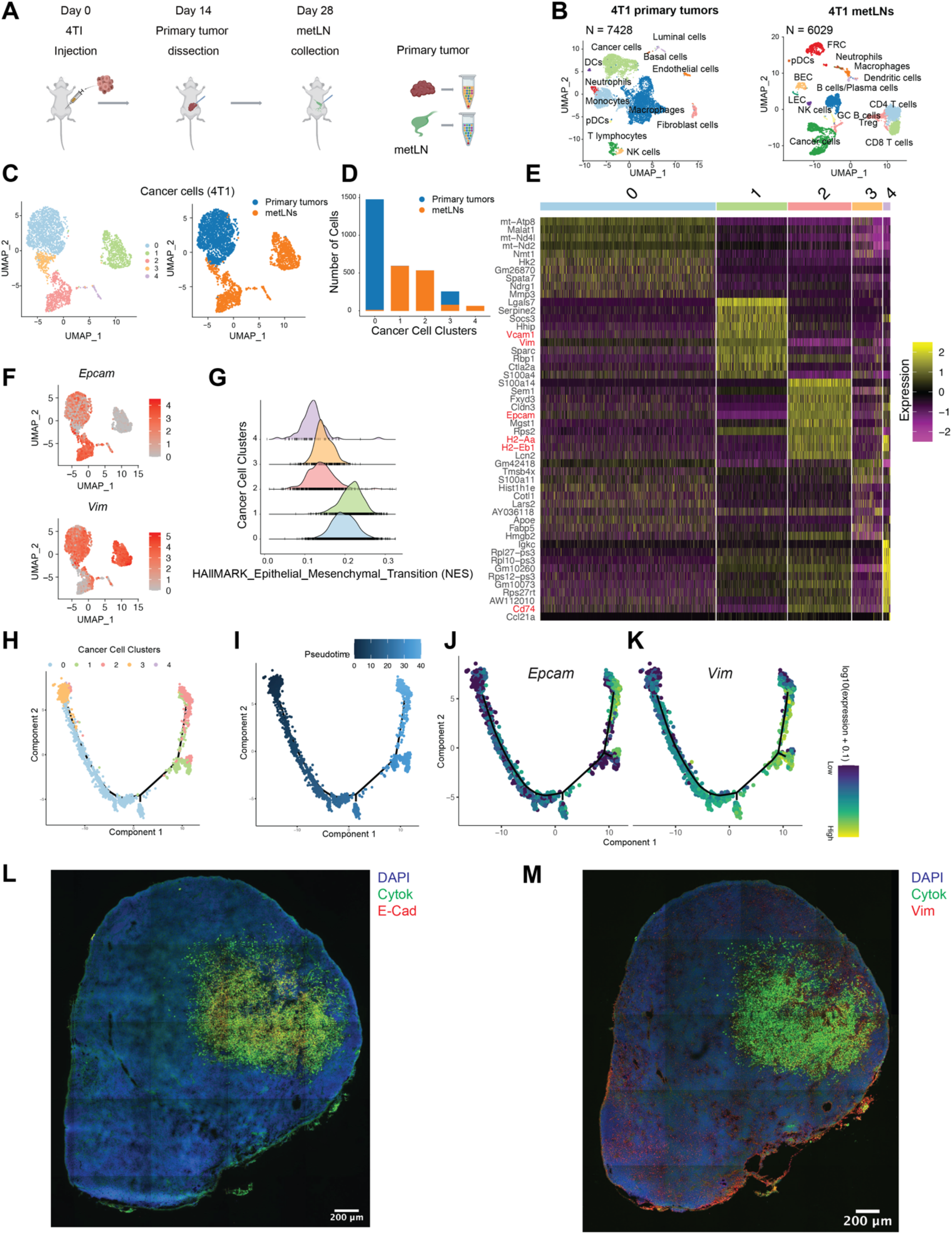
Single-cell RNASeq identified EMT-plasticity of cancer cells in 4T1 primary tumors and metastatic lymph nodes. **A)** Pipeline for single-cell RNA sequencing library preparation. Briefly, we injected 1×10^5^ 4T1 cells into the 4^th^ mammary fat pad of 6∼10-week-old female Balb/c mice. On day 14, or when tumor size reached 500 mm^3^, the primary tumors were dissected and dissociated into single cells for 10X Genomics Chromium Single Cell 3’ library preparation. Fourteen days after primary tumors resection, the tumor metastatic lymph nodes (metLNs) were collected, digested into single cells and processed by the 10x Genomics platform. Illumina NextSeq platform employed for sequencing. **B)** UMAP of distinct types of cells in 4T1 primary tumors (left) and metLNs (right) microenvironment. **C)** The UMAP of aggregated cancer cells (n = 3007) from primary tumors and metLNs after cell cycle regression, (left) colored by original cluster names, (right) colored by samples. Top 20 principal components were chosen for the UMAP analysis, minimum distance is 0.5. **D)** The numbers of cancer cells in each cluster in panel **c. E)** Heatmap of single-cell gene expression of the top 10 differentially expressed genes in the cancer cells. The color bar above the heatmap is the same color code as panel **c**. In the heatmap, purple represents low expression, and yellow represents high expression. **F)** UMAP of gene expression of epithelial cell marker genes (*EpCAM*) and mesenchymal cell marker genes (*Vim*) in cancer cells. Gene expression values are log normalized, gray represents low expression, and red represents high expression. **G)** Histogram of the single-cell enrichment score of MSigDB Hallmark Epithelial Mesenchymal Transition gene set. **H)** The single-cell trajectory of cancer cells colored by original cluster identities. **I)** The pseudotime trajectory of cancer cells, dark color represents early stage, bright color represents late stage. **J-K)** Expression of *EpCAM* and *Vim* were projected to the single-cell trajectories. Gene expression values are scaled, and log normalized. **L-M)** Representative confocal microscopy images of immunofluorescence staining of cytokeratin, E-cadherin and vimentin in 4T1 metLN. Two adjacent tissue sections from the same lymph node were used for the staining. The thickness of the tissue section is 10 μm. cytokeratin (1:200 diluted), E-cadherin (1:100 diluted), vimentin (1:100 diluted). Scale bars as marked on figure.

### Lineage analysis of epithelial to mesenchymal transition (EMT) remodeling during breast cancer progression to lymph node metastasis

To determine the intra-tumoral gene-expression heterogeneity during lymph node metastasis, we integrated cancer cells from primary tumors and metLNs and analyzed them by Uniform Manifold Approximation and Projection (UMAP) (**Figure 1C**). The cancer cells were grouped into 5 different clusters, and interestingly cancer cells in primary tumors metLNs are clearly separated from each other. Further, cancer cells from the metLNs segregated into two distinct major populations (cluster1 and cluster2) (**Figure 1D**). Next, we examined the cancer cell heterogeneity by comparing differentially expressed genes of the 5 cancer cell clusters. The top 10 most significantly differentially expressed marker genes demonstrate that *Vcam1* and *Vim*, the mesenchymal cell marker genes, were highly expressed in cluster1 cancer cells from metLNs. In contrast, *Epcam*, a marker gene for epithelial cells, was highly expressed in clusters2 and cluster4 cancer cells from metLNs (**Figure 1E**). We further projected the single cell gene expression of *Epcam* and *Vim* into the cancer cell UMAP. The cancer cells in primary tumors expressed both *Epcam* and *Vim* (cluster0 and part of cluster3), suggesting these are the EMT-hybrid cells. While cancer cells in metLNs were either *Epcam* positive or *Vim* positive, with a small population of *Epcam* and *Vim* double positive cells (**Figure 1F**). Next, we also examined the single cell gene set enrichment score of the cancer cells based on the MSigDB Hallmark genesets. We found that the cancer cells in cluster0 and cluster1 had a higher epithelial mesenchymal transition score, with cancer cells in cluster1 exhibiting the highest enrichment score of the epithelial to mesenchymal transition signature. The cluster2, 3, 4 cancer cells showed lower levels of EMT signature, indicating a more epithelial-like phenotype (**Figure 1G**). Surprisingly, *Cd74, H2-Eb1* and *H2-Aa*, key components of the MHC-II receptor complex needed for antigen-presentation, were highly expressed in the epithelial-like cancer cells (**Figure 1E**).

Increasing evidence suggests EMT status is a ‘spectrum’ rather than a binary status^26^. Furthermore, this transition is reversible as mesenchymal-like cells can also transition to an epithelial phenotype (MET)^27^. To investigate EMT/MET during lymph node metastasis, we examined the lineage of cancer cells during lymph node metastasis by single-cell trajectory analysis^28,29^. We found that most of the cancer cells in cluster2 were in the late stage of the pseudotime trajectory (**Figure 1H-I**). Next, we projected the gene expression of *Epcam* and *Vim* into the pseudotime trajectory. Of note, we found that epithelial cancer cells in the metLNs were derived from mesenchymal cancer cells (**Figure 1J-K)**, suggesting MET of lymph node metastasis cancer cells.

We further investigated the EMT phenotype of cancer cells and the metastatic burden in individual lymph nodes by flow cytometry (**Figure S2A**). We found that in the metLNs in which cancer cells account for more than 5% of the cell population, nearly 80% of the cancer cells were EpCAM positive. In contrast, in the metLNs in which cancer cells account for less than 1% of the cell population, on average around 14% of the cancer cells were mesenchymal phenotype (EpCAM-vimentin+) and 19% of the cancer cells were epithelial phenotype (EpCAM+vimentin-) (**Figure S2B-C**). Using immunostaining of EMT hallmarks in metLNs, we found both E-cadherin positive (epithelial) cancer cells and vimentin positive (mesenchymal) cancer cells in LNMs, with most of the E-cadherin positive cancer cells in the center of the metastatic lesion (**Figure 1L-M**). These data demonstrate that the metastatic cancer cells in metLNs displayed heterogenous and spatially organized epithelial and mesenchymal phenotypes in metastatic lesions.

Taken together, these results indicate that cancer cells in the primary tumor underwent EMT, which enhanced the ability of the more mesenchymal cancer cells to metastasize to the lymph node. However, in the metLN, a portion of the mesenchymal cancer cells underwent MET, particularly those in the center of the lesion where they are less exposed to the native lymph node microenvironment.

### Dynamic transcriptomic alterations during breast cancer lymph node metastasis

EMT/MET is a common phenotype during cancer progression and is an attractive target in the clinic. However, direct targeting of EMT molecules remains a challenge. Understanding how EMT/MET reprograms cancer cells to adapt to the lymph node microenvironment could potentially identify new signaling pathways involved in EMT and provide novel strategies to indirectly target this process. We ranked cancer cells based on the pseudotime trajectory and compared the differentially expressed gene signatures among them. Besides the elevation of EMT and MET gene signatures (**Figure S2D**), we also found that type I and II interferon signaling, fatty acid metabolism and IL6/JAK/STAT3 signaling were elevated in cancer cells during the progression of breast cancer to lymph node metastasis (**Figure S2E**).

The elevation of IL6/JAK/STAT3 signaling pathway in cancer cells induces a more invasive and advanced phenotype in breast cancer, ovarian cancer and prostate cancer^30–32^. IFN-γ and JAK/STAT induced gene activation accounts for the induction of MHC-I and MHC-II on antigen-presenting cells^33^. Published data suggest that EMT processes protect mesenchymal cancer cells from immune surveillance in the primary tumor by down-regulation of MHC-I molecules^34^. However, in our data, we found that MHC-I molecules were generally highly expressed in all cancer cells (**Figure S3A**). In contrast, MHC-II molecules were expressed in a fraction of primary cancer cells and were robustly induced during the progression of breast cancer to lymph node metastasis, particularly in epithelial-like metastatic cancer cells (cluster2 and cluster4) (**Figure 2A**). Single-cell pseudotime trajectory results suggest the elevation in gene-expression of MHC-II molecules (*H2-Aa* and *H2-Ab1*) on late-stage cancer cells that have undergone MET (**Figure 2B-C**). Although MHC-II is predominantly displayed on professional antigen-presenting cells (APCs), we and others have recently demonstrated expression and functional significance of MHC-II in both malignant^32,35,36^ and normal epithelial^37,38^ as well as lymphatic endothelial cells^39^. CD4+ T cell recognition of cognate antigen bound to MHC-II in the absence of co-stimulation leads to T cell tolerance through anergy^36^ or induction of regulatory T cell (Treg) differentiation^40^. Accordingly, we found that the LNM cancer cells lacked expression of co-stimulatory molecules *Cd80, Cd86*, and *Icosl* (**Figure 2D**). The elevation of type II interferon signaling pathway in cancer cells has been shown to mediate the expression of co-inhibitory molecule PD-L1 to enhance immune evasion^41^. However, the expression of *Cd274*—the gene that encodes PD-L1—was almost undetectable in LNM cancer cells, suggesting cancer cells might rely on PD-L1 independent strategies to escape immune surveillance in the lymph node.

**Figure2:**
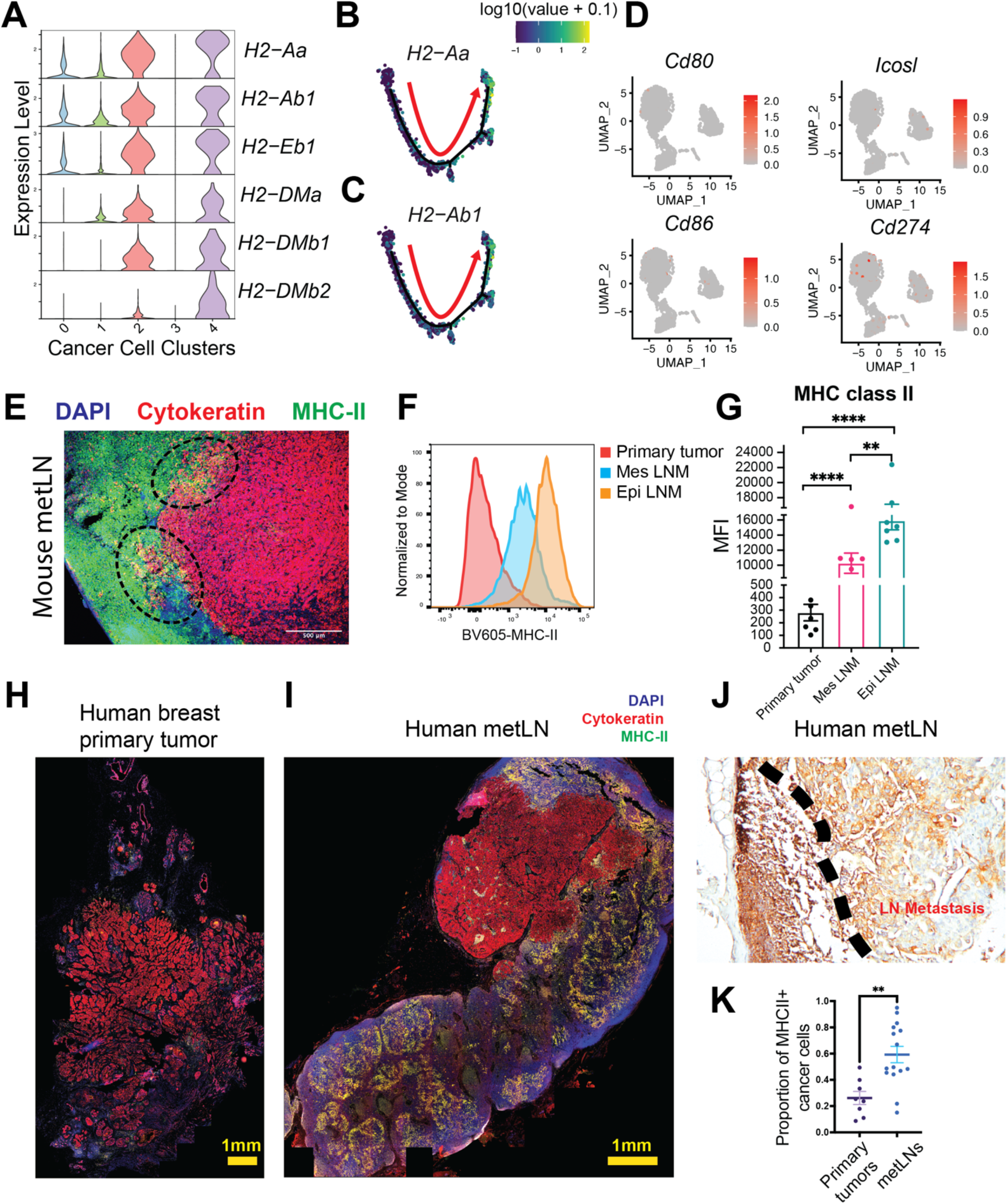
The presence of MHC-II on cancer cells during lymph node metastasis. **A)** Violin plot represents the gene expression of MHC-II key molecules in cancer cells (clusters 1, 2, 4 from metLNs). **B-C**) Expression of MHC-II key molecules *H2-Aa* and *H2-Ab1* were projected to the single-cell trajectories. Gene expression values are scaled, and log normalized. The red arrow indicates the pseudotime trajectory of cancer cells progression. **D)** UMAP of single cell gene expression of co-stimulatory (*CD80, CD86, Icosl*) and co-inhibitory (*CD274*) molecules in cancer cells. **E)** Immunofluorescence staining of MHC-II-expressing 4T1 cancer cells in murine metLNs. Representative confocal microscopy images of nuclei (DAPI, blue), pan-cytokeratin (red) and MHC-II (green). The thickness of the tissue section is 10 μm. **F)** Cancer cell surface MHC-II intensity measured by flow cytometry. **G)** The mean fluorescent intensity (MFI) of MHC-II on cancer cells from 4T1 primary tumors and metLNs. Mes LNM: EpCAM-vimentin+; Epi LNM: EpCAM+vimentin-. One-way ANOVA was used for the statistical analysis. ** p-value < 0.01, **** p-value < 0.0001. Tukey’s multiple comparisons test was used for the post hoc test. **H-I)** Immunofluorescence staining of nuclei (DAPI, blue), pan-cytokeratin (red) and MHC-II (green) in human breast cancer primary tumor (**H**) and matched metastatic lymph node (**I**). The thickness of the tissue section is 5 μm. **J)** Immunohistochemistry staining of MHC-II (HLA-D) in human breast cancer metastatic lymph node. **K)** The proportion of MHC-II positive cancer cells in human breast tumors (n=8) and lymph node metastases (n=15). ** P-value < 0.01. Student’s t-test was used for the statistical analysis.

We verified the presence of MHC-II on cancer cells in metLNs via immunofluorescence staining of 4T1 metLNs (**Figure 2E**). We further examined cell surface MHC-II in primary tumor and LNM cancer cells via flow cytometry (**Figure 2F**). We found that EpCAM+/vimentin– (epithelial) cancer cells in metLNs exhibited the highest MHC-II on their cell surface (**Figure 2F-G**). Consistent with these findings, we also found the presence of MHC-II+ cancer cells in metLNs from patients with invasive ductal breast carcinoma (**Figure 2H-J**). The proportion of MHC-II+ cancer cells in metLNs were significantly (p-value < 0.01) higher than in the primary tumors in these patient samples (**Figure 2K**). In some of these patients, these MHC-II+ cancer cells displayed a more invasive phenotype (**Figure S3**), which is similar to what was observed in 4T1 metLNs (**Figure 2E**). All together, these results suggest that breast cancer cells upregulate MHC-II without concurrent expression of co-stimulatory molecules during the progression of breast cancer to lymph node metastasis in both mice and humans. We hypothesized that cancer cell MHC-II expression likely contributes to immune evasion by eliciting T cell tolerance.

### IFN-γ signaling pathway induces MHC class II expression in cancer cells

IFN-γ has been shown to regulate MHC class I/II expression in epithelial cells via the JAK/STAT signaling pathway^33,35,37^. The elevation of the interferon gamma response gene signature (**Figure 3A**) and the increase in expression of IFN-γ receptors *Ifngr1/2* and *Ciita*, the transactivator of MHC-II, during breast cancer progression to lymph node metastasis (**Figure S4A-C**) led us to hypothesize that this pathway is involved in the elevation of MHC-II on LNM cancer cells. *In vitro* administration of IFN-γ to 4T1 cells led to a profound increase in the mRNA expression of the key MHC-II molecules *H2-Aa* and *H2-Ab1* (**Figure 3B**). Of note, we also observed a significant (*p*-value < 0.001) increase in MHC-II+ cells and cell surface MHC-II molecules post IFN-γ stimulation compared to control group (**Figure 3C-D**). Further, we found that IFN-γ was able to induce mRNA expression of MHC-II genes (*H2-Aa, H2-Ab1* and *H2-DMa*) in additional cancer models B16F10 (melanoma), E0771 (breast) and MCa-P1362 (breast)^19^ (**Figure S4D-F**). The MHC-II transactivator gene CIITA is a general regulator of both inducible and constitutive MHC-II expression^33^. To understand whether the IFN-γ induced expression of MHC-II on cancer cells is CIITA dependent, we generated the *Ifngr1/2* and *Ciita* knock-out 4T1 cells using CRISPR/Cas9 systems. In *Ifngr1/2* and *Ciita* knock-out cells, the administration of IFN-γ was not able to induce mRNA expression of *H2-Aa* and *H2-Ab1* (**Figure 3E**) and cell surface MHC-II (**Figure 3F**), confirming that IFN-γ signaling pathway and CIITA are essential for the MHC-II expression on 4T1 cancer cells.

**Figure3:**
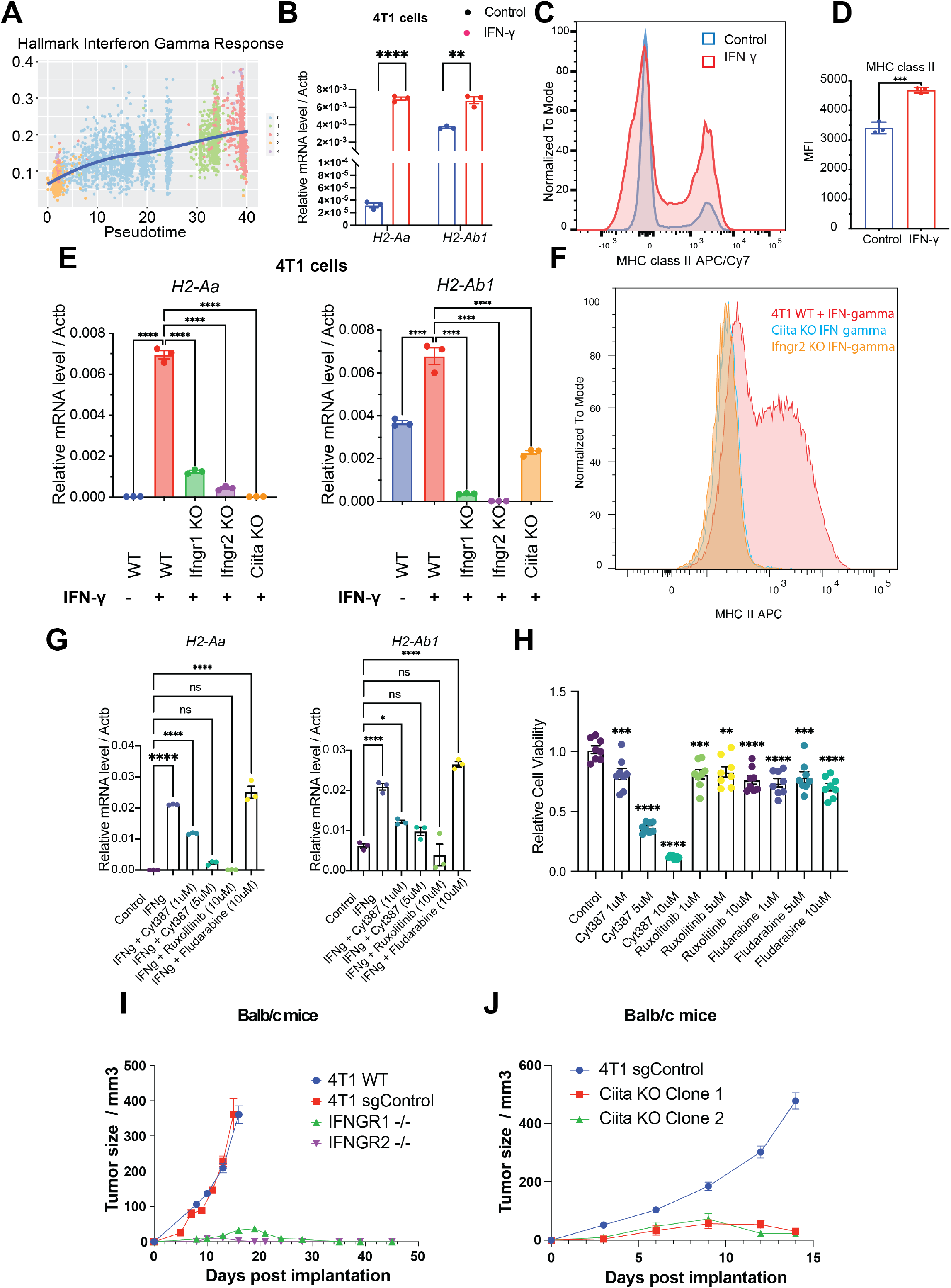
The induction of interferon gamma signaling pathway and MHC-II are critical for tumor progression. **A)** The single-cell enrichment score of the MSigDB Hallmark Interferon Gamma Response gene set. Cancer cells were ranked by pseudotime, and the blue line represents the loess regression of the enrichment score. **B)** IFN-γ induced expression of MHC-II molecules *H2-Aa* and *H2-Ab1 in vitro*. 4T1 cells were treated with or without IFN-γ (10 ng/mL) for 24 hours. Student’s t test for statistical analysis, ** p-value < 0.01, **** p-value < 0.0001. **C)** IFN-γ induced cell surface MHC-II on 4T1 cells. 4T1 cells were treated with or without IFN-γ (10 ng/mL) for 24 hours and cell surface MHC-II was measured by flow cytometry. **D)** The mean fluorescent intensity (MFI) of MHC-II on MHC-II positive 4T1 cancer cells. **E)** Expression of MHC-II molecules *H2-Aa* and *H2-Ab1* in 4T1 WT, *Ifngr1, Ifngr2 and Ciita* knock-out cells that with or without IFN-γ (10 ng/mL) treatment for 24 hours. One-way ANOVA was used for the statistical analysis, ** p-value < 0.01, *** p-value < 0.001, **** p-value < 0.0001. Tukey’s multiple comparisons test was used for the post hoc test. **F)** The cell surface MHC-II on 4T1 WT, *Ifngr2 and Ciita* knock-out cells post IFN-γ (10 ng/mL) treatment for 24 hours. **G**) Expression of MHC-II molecules *H2-Aa* and *H2-Ab1* mRNA in 4T1 cells with IFN-γ or IFN-γ combined with JAK/STAT inhibitor treatment. One-way ANOVA was used for the statistical analysis, ** p-value < 0.01, *** p-value < 0.001, **** p-value < 0.0001. Tukey’s multiple comparisons test was used for the post hoc test. **H)** Cell viability under different JAK/STAT inhibitor treatment. 4T1 cells were treated with DMSO, Cyt387, Ruxolitinib, and Fludarabine for 72 hours. Cell viability was measured by MTT assay. **I)** The tumor growth of 4T1 WT (N=4), 4T1 sgRNA control (N=4), and *Ifngr1/2* knockout cancer cells (N=12) in Balb/c mice. **J)** The tumor growth of 4T1 sgRNA control and *Ciita* knockout cancer cells in Balb/c mice (N=4). Two different *Ciita* knockout cell clones were selected for the *in vivo* assay.

We previously showed that downstream of IFN-γ signaling, the JAK/STAT pathway mediates the induction of epithelial MHC-II expression in the small intestine^37^. As the IL6/JAK/STAT signaling pathway was upregulated in LNM cancer cells (**Figure S2E**), we tested whether targeting the JAK/STAT pathway is capable of suppressing MHC-II expression. Administration of JAK/STAT inhibitors Cyt387 (momelotinib), ruxolitinib and fludarabine in combination with IFN-γ to 4T1 cancer cells for 48 hours showed that JAK1/2 inhibitors Cyt387 and Ruxolitinib can suppress IFN-γ induced MHC-II mRNA expression on 4T1 cancer cells (**Figure 3G**). Together, these results suggest that MHC-II expression in breast cancer cells is regulated by IFN-γ induced JAK/STAT signaling.

We next tested whether pharmacologic inhibition of JAK/STAT signaling influences tumor progression in the 4T1 orthotopic breast cancer model. Treatment of 4T1 cancer cells with a JAK1/2 inhibitor Cyt387 showed direct anti-cancer cell activity at 5mM and 10mM concentration *in vitro*, whereas ruxolitinib and fludarabine did not (**Figure 3H**). Next, we injected 100,000 4T1 cancer cells into the mammary fat pad of Balb/c mice, and at day 7 we started the treatment with JAK/STAT inhibitor Cyt387 every two days for 7 doses (**Figure S4G**). The growth of tumor (**Figure S4H**) and weight of mice (**Figure S4I**) did not show a significant difference between the control and Cyt387 treated group. The draining inguinal lymph nodes in mice treated with Cyt387 were smaller compared with the lymph nodes in the control group (**Figure S4J**), and the Cyt387 treated group had more metastatic nodules in the lung (**Figure S4K**). These results could be due to the fact that Cyt387 also will inhibit JAK/STAT signaling in immune cells, which are necessary for anti-tumor immune responses.

To test if cancer cell-intrinsic IFN-γ signaling-mediated activation of CIITA and concomitant elevation of MHC-II on cancer cells influence breast cancer tumor progression, we orthotopically implanted the *Ifngr1/2* or *Ciita* knockout cells as well as the sgRNA control cells into the mammary fat pad of wild type Balb/c mice. There was no difference in tumor growth between 4T1 wild type cells and sgRNA control cells (**Figure 3I**). However, ablation of *Ifngr1/2* or *Ciita* in cancer cells significantly prevented breast cancer progression in the wild type Balb/c mice (**Figure 3I-J**), but not in immunodeficient mice (**Figure S4L-M**), suggesting that IFNGR-CIITA axis is a critical mediator of immune evasion during breast cancer progression. These results are concordant with a previous study that demonstrated that a lack of *Ifngr2* and *Jak1* expression in melanoma cells blunts tumor growth in immunocompetent mice^42^. Taken together, our results show the critical role of the IFNGR-JAK/STAT-CIITA signaling pathway axis in regulating MHC-II expression on cancer cells and immune evasion during breast cancer progression.

### Invasion of cancer cells in metLNs enhances Treg activation and elicits an immunosuppressive microenvironment

Induction of T cell anergy is a key mechanism of self-tolerance. Anergic T cells can also further expand the CD4+Foxp3+ Treg repertoire^43^. To investigate whether MHC-II on epithelial-like cancer cells leads to CD4+ T cell tolerance and enhances the expansion of Tregs in metLNs, we investigated the MHC-II signaling pathway cell-cell interaction network in primary tumors and metLNs^44^. Cancer cells in the primary tumor interacted with pDCs more frequently (**Figure 4A**), while cancer cells in the metLNs interacted with CD4+ T cells and Treg cells as well as pDCs (**Figure 4B**). The ligand-receptor pair analysis predicted that MHC-II molecules on cancer cells interact with CD4 and LAG3 on CD4+ T cells, Treg and pDCs in metLNs (**Figure 4C**). To understand the impact of cancer cell invasion on the lymph node microenvironment, we performed single-cell RNA sequencing in the inguinal lymph nodes from naïve Balb/c mice (**Figure S5A-C**). We aggregated single-cell mRNA expression data from naïve lymph nodes with non-cancer cells from metLNs by UMAP analysis to be able to compare expression profiles of specific T cell populations from naïve lymph nodes and metLNs (**Figure S5D-F**). Gene ontology functional annotation of the differentially expressed genes in Treg cells between metLN and naïve LN demonstrated that Aurora A/B signaling, PLK1 signaling, proliferation-associated transcription factor FOXM1 network were up-regulated in Tregs in metLNs, suggesting they were highly proliferative (**Figure 4D**). Additionally, androgen receptor signaling and TGF-β receptor signaling pathways were also upregulated in Tregs in metLNs (**Figure 4D**). The T cell intrinsic androgen receptor activity limits anti-tumor immunity and T cell re-invigoration^45,46^. TGF-β induces Foxp3+ Treg cells from CD4+CD25-precursors^47^ and helps maintaining Treg suppressor function^48^. Thus, our data suggests that in the metLNs Tregs exhibited a more immunosuppressive phenotype compared to naïve LN Tregs. Furthermore, we found that Tregs in both naïve lymph nodes and metLNs expressed the inhibitory checkpoint molecule *Ctla4*, but not *Pdcd1* (PD-1) (**Figure 4E**). The co-stimulatory molecules *Tnfrsf18* (GITR) and *Tnfrsf4* (OX-40) were prevalently expressed in Tregs, however only the Tregs in metLNs expressed *Icos* (**Figure 4E**). Moreover, Treg activation associated genes and T cell anergy and exhaustion associated molecules, such as *Foxp3, Il2ra* (CD25), *Cd44, Ikzf2* (Helios*), Maf, Tox* and *Izumo1r* (FR4), were profoundly elevated in Tregs in metLNs (**Figure 4E**), suggesting enhanced immunosuppressive characteristics^49,50^.

**Figure4:**
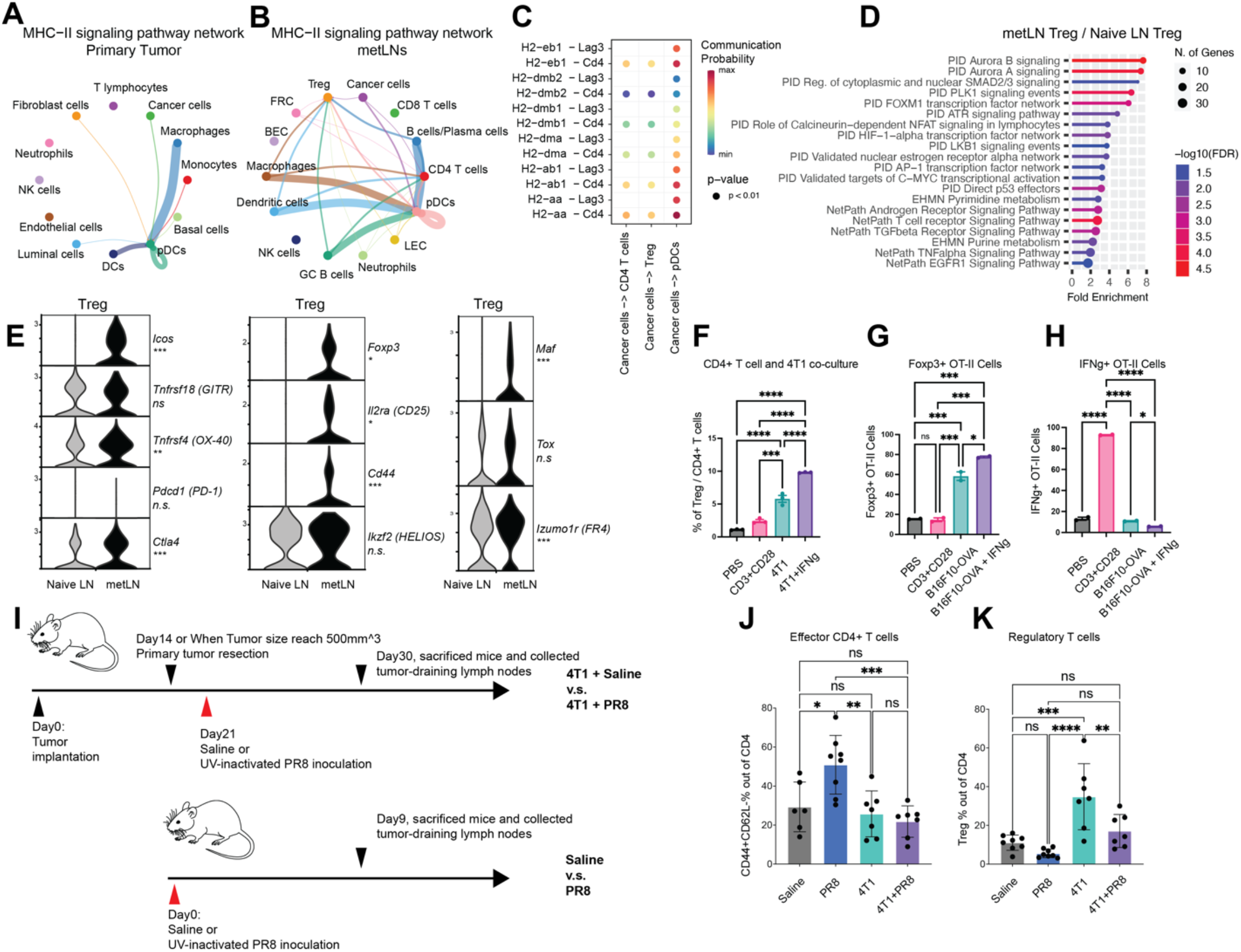
MHC-II positive cancer cells mediated the enrichment of tolerogenic Treg cells in lymph nodes through anergy. **A-B)** Single-cell MHC-II signaling pathway network in (**A**) 4T1 primary tumor and (**B**) metLNs. The cell-cell interaction analysis was performed by CellChat. **C)** MHC-II signaling pathway ligand-receptor interactions between cancer cells and CD4+ T cells, Treg, and pDCs in metLNs. **D)** Functional annotation of up-regulated genes in Tregs in metLNs by ShinyGO CuratedGeneSetDB. **E**) The single-cell gene expression of Treg cell associated genes. **F)** The percentage of regulatory T cells when cultured *in* vitro with or without 4T1 cancer cells. 4T1 cells were pretreated with or without IFN-γ (10ng/mL) for 48 hours before the coculture assay. **G)** The percentage of Foxp3+ OT-II T cells in the *in vitro* co-culture assay. **H)** The percentage of IFN-γ+ OT-II T cells in the *in vitro* co-culture assay. B16F10-OVA cells were pretreated with or without IFN-γ (10ng/mL) for 48 hours before the coculture assay. **I)** The experiment design of UV-inactivated influenza virus PR8 inoculation assay. **J-K)** The percentage of (J) effector CD4+ T cells and (K) regulatory T cells in naive or metastatic lymph nodes with or without inoculation with UV-inactivated influenza virus PR8.

Next, we tested whether the induction of MHC-II on 4T1 cancer cells in the absence of co-stimulatory signals can lead to CD4+ T cell anergy and expansion of Tregs. We treated 4T1 cancer cells with or without IFN-γ for 48 hours, then isolated CD4+ T cells from 12-week-old female Balb/c mice and cocultured them *in vitro* for 72 hours. PBS and anti-CD3/anti-CD28 coated wells served as T cell activation negative and positive controls, respectively. When CD4+ T cells were cultured alone in the presence of both anti-CD3 (signal 1) and anti-CD28 (signal 2), we observed around 2% of Treg cells. In contrast, we observed 2-fold increase of the percentage of Treg cells when co-cultured with 4T1 cells and 4-fold increase of percentage of Treg cells in the presence of IFN-γ pretreated 4T1 cells (MHC-II+) compared to the anti-CD3+CD28 group monoculture group (**Figure 4F**). We further examined the interaction between cancer cell MHC-II and TCRs on CD4+ T cells with an antigen-specific B16F10-OVA/OT-II coculture assay. Very similarly, in the presence of IFN-γ pretreated B16F10-OVA cancer cells, there was 3∼4-fold increase of Foxp3+ OT-II cells compare to the anti-CD3+CD28 group (**Figure 4G**). In addition, we observed the lowest population of IFN-γ + OT-II cells in the presence of IFN-γ pretreated B16F10-OVA cancer cells (**Figure 4H**).

To further evaluate the immune consequences of tumor metastasis in the draining lymph nodes *in vivo*, I tested immune responses of metLNs against UV-inactivated PR8 viruses, a well-recognized model to study the immune activation in the lymph nodes^51,52^. We compared the T cell phenotype in normal lymph nodes, UV-inactivated PR8 influenza virus inoculated lymph, and 4T1 metLNs as well as 4T1 metLNs inoculated with UV-inactivated PR8 influenza virus (**Figure 4I**). We found that the population of effector CD4+ T cells remained similar between normal lymph nodes and metLNs. However, the immune activated lymph nodes had higher effector CD4+ T cells (**Figure 4J**). In contrast, in metLNs inoculated with UV-inactivated PR8 influenza virus, we did not see an increase in effector CD4+ T cells. Further, there was a substantial increase in Tregs in 4T1 metLNs (**Figure 4K**). Taken together, these results suggest that the invasion of cancer cells into the lymph node triggers the generation and activation of Tregs, leading to an immunosuppressive microenvironment of the metLNs.

### Invasion of cancer cells remodels human breast cancer sentinel lymph node microenvironment

To understand the clinical significance of our findings, we utilized a human breast cancer single-cell transcriptome dataset with paired primary tumors and axillary lymph nodes from the same side^53^. This dataset consists of specimens from five patients, including 1 Luminal B breast cancer patient, 2 Her2+ breast cancer patients and 2 triple negative breast cancer (TNBC) patients. For each of these patients, there were 2 lymph nodes collected for single-cell sequencing, including one metastasis positive lymph node and one cancer negative tumor-draining lymph node.

In the breast cancer primary tumors, we characterized 27593 single-cells that group into fibroblasts, cancer cells, myeloid cells, T cells, plasmablasts, endothelial cells, B cells and pericytes (**Figure S6A-B**). In the metLNs, we characterized 25739 single-cells and identified B cells, T cells, cancer cells, myeloid cells, plasmablasts, FRCs and LECs based on their marker genes (**Figure S6C-D**). We integrated only the cancer cells from primary tumors and metLNs from patients with lymph node metastasis. Interestingly, we found that cancer cells from the same patient cluster closer to each other (**Figure 5A**), showing the expected inter-patient cancer cell heterogeneity. Next, we examined the expression of MHC-II molecules and gene signatures in all cancer cells. We found that both Her2+ and TNBC breast cancer LNMs showed expression of MHC-II molecules and their MHC-II signatures were higher compared to primary tumors (**Figure 5B-C**). In contrast, the MHC-I gene signature and expression of MHC-I molecules were reduced in metLNs compared to the primary tumor in these human data (**Figure S6E-F**) These results suggest the upregulation of MHC-II is a more universal phenotype in breast cancer that can lead to cancer cell immune evasion in lymph nodes.

**Figure5.**
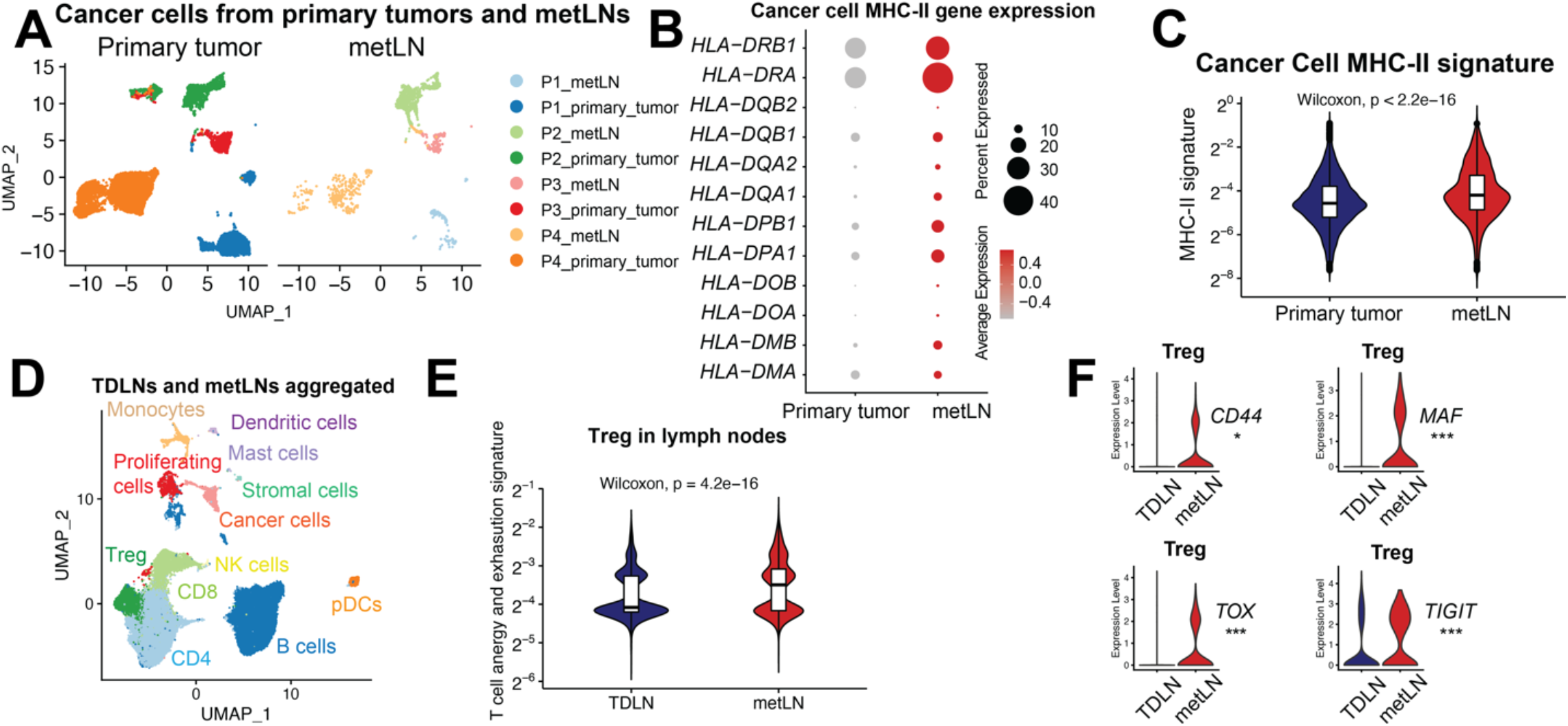
Dynamic change of T cell states in human breast cancer metastatic lymph nodes. **A)** The UMAP of 14646 aggregated cancer cells from primary breast tumor and metLNs. The aggregated cancer cell UMAP is separated to primary tumor (left) and metLN (right) for visualization. We selected the top 30 principal components for the UMAP analysis with a minimum distance of 0.5 for the display. **B**) The expression profiling of MHC-II genes in cancer cells from primary tumors and metLNs. Red represents high expression; gray represents low expression. The size of the circle represents the proportion of cells expressing the indicated genes in each cluster. **C**) The MHC-II gene signature in cancer cells from primary tumors and metLNs. Single-cell MHC-II signature score was measured by UCell (https://github.com/carmonalab/UCell. **D**) The UMAP of cells from TDLNs and metLNs colored by cell types. **E**) Violin plot shows the anergy and exhaustion gene signature in Tregs in TDLNs and metLNs from all the patients. Gene expression signature score was measured by UCell. The CD4+ T cell anergy and exhaustion genes were generated by Trefzer et al., 2021, Cell Reports. **F**) Violin plots show the gene expression of *CD44, MAF, TOX*, and *TIGIT* in Treg cells in the lymph nodes. Wilcoxon Rank-Sum test was used for the statistical analysis.

To understand the impact of cancer cell invasion on T cells in patient lymph nodes, we integrated the lymph node single-cell datasets from patients with lymph node metastasis to be able to compare the T cells phenotypes between TDLNs and metLNs (**Figure 5D**). Using single cell RNA sequencing data, we identified all the T cells and Tregs from patient lymph nodes using cell-specific gene signatures. Using a regulatable antigen presentation system, a recent study characterized a comprehensive CD4+ T cell anergy and exhaustion gene signature after exposure to persistent antigen stimulation^50^. Using this CD4+ T cell anergy and exhaustion gene signature as a reference, we found that Tregs cells in TDLN and metLN from the same patient exhibited very different phenotype. In metLNs, the Tregs cells showed increasing of T cell anergy and exhaustion gene signatures (**Figure 5E**), indicating enhanced immunosuppressive characteristics^49,54,55^. Of note, in one of the patients who had the largest numbers of cancer cells in their metLN (**Figure 5A**), we found the elevation of *CD44, TOX, MAF* and *TIGIT* in Treg cells in metLNs (**Figure 5F**), which is constitent with metLNs in the mouse models (**Figure 4E**). These data indicate the presence of MHC-II on cancer cells in metLNs correlates with an immunosuppressive lymph node microenvironment.

Altogether, our study shows that in the progression of breast cancer cells to the lymph node, metastatic breast cancer cells displayed significant plasticity. Further, a subset of LNM cancer cells upregulate MHC-II in metLNs to induce immune tolerance and evade anti-tumor immunity.

## Discussion

Many epithelial-origin cancer cells undergo EMT during metastasis^56–60^. EMT reprograms expression of tight junction and cell membrane proteins, with a decrease of E-cadherin and epithelial cell adhesion molecule (EpCAM) as well as an increase of N-cadherin and vimentin^61–63^. These changes facilitate cancer cell detachment and migration, which are key steps in dissemination. Studies have demonstrated that EMT of cancer cells in the primary tumor results in increased lymph node metastases^64^. By using single-cell RNA sequencing in human triple-negative breast tumors, a recent study showed mixed populations of malignant cells and various states of cancer cell EMT^56^. In our data, most of the cancer cells in 4T1 primary tumors were *Krt8, Krt18* and *EpCAM* triple positive. The mesenchymal cell marker genes, *Twist1* and *Vim*, were also expressed in the *EpCAM* positive cancer cells, revealing these cells were in an EMT-hybrid state^65^. Recent studies have shown that the transition of fully epithelial cells into fully mesenchymal cells occurs gradually^26^, increasing the probability of identifying cells along a continuum of the EMT process when looking at a single snapshot in time. Cancer cells in an EMT hybrid state exhibited increased invasion and tumor propagation^66^.

We and others observed that metastatic cancer cells enter into the lymph node from the subcapsular sinus, and then invade into the cortex and medullary areas^67,68^. Furthermore, we and others have shown that metastatic cancer cells can exit the lymph node through lymph node blood vessels and spread to distant sites^10,11^. In metLNs, we found both mesenchymal-like and epithelial-like cancer cells (**Figure 1F, L, M**). Most of the epithelial-like cancer cells were round and remained closely attached to one another in the center of LNM lesions, although some remained at the tumor margin (**Figure 1L**). In contrast, the mesenchymal-like cancer cells were spindle-shaped and located around the margin of LNM lesions, close to the lymph node subcapsular sinus (**Figure 1M**). Supported by our data, we hypothesize that the LNM mesenchymal cancer cells are ‘seeds’ from the primary tumor, some of which undergo MET and differentiate back to an epithelial phenotype in the center of the lesion where they are shielded from the native lymph node microenvironment and can better recreate a microenvironment that resembles the primary tumor. Supporting this concept, single-cell trajectory analysis revealed that LNM epithelial cancer cells were derived from LNM mesenchymal cancer cells in lymph nodes (**Figure 1H-K**).

In addition to cancer cell plasticity during breast cancer progression to lymph node metastasis, we found that the genes encoding MHC-II molecules were elevated in LNM epithelial cancer cells (**Figure 2**). The MHC-II complex is critical to the presentation of antigens to CD4+ T cells and is predominantly present on professional antigen-presenting cells. However, epithelial cells—which are normally well-positioned and covering all body surfaces—can also express MHC-II molecules and act as non-professional antigen-presenting cells^36–38^, helping to preserve immune tolerance or surveillance at mucosal sites^69^. Intriguingly, MHC-II+ LNM cancer cells lacked co-stimulatory molecules, including *CD80* and *CD86* (**Figure 2D**), which are essential for the activation of CD4+ T cells. In the absence of co-stimulation, MHC-II presence can promote tolerance through T cell anergy or induction of Treg differentiation^40^. Consistent with this, we found high expression of *Foxp3, Il2ra* (CD25), *Icos, Tnfrsf18* (GITR), *Maf, Tox* and *Ikzf2* (Helios) in Tregs from the metLNs (**Figure 4E**). *In vitro* co-culture of T cells with 4T1 cancer cells also showed a decrease of IFN-γ+ CD4+ T cells and an increase of Tregs (**Figure 4F-H**). Further, we measured impaired CD4+ T cell activation and greater numbers of Tregs in metLNs (**Figure 4J-K**).

Of note, we also found the presence of MHC-II+ cancer cells in human breast cancer metLNs (**Figure 2I-J and 5B**), highlighting the potential clinical significance of our findings. Cancer cells upregulated MHC-II across their progression trajectory from primary tumor to lymph node metastasis. Interestingly, the MHC-II+ cancer cells present a more invasive characteristic, in both murine (**Figure 2E**) and human breast cancer metLNs (**Figure S3B**). The invasion and presence of MHC-II+ cancer cells in lymph nodes showed a positive correlation with more immunosuppressive Treg cells (**Figure 4E, 5E-F**). Notably, ablation of *Ifngr1/2* or *Ciita* blunted cancer-intrinsic MHC-II expression and prevented tumor progression. Future studies are needed to delineate the precise kinetics of cancer-intrinsic MHC-II expression, cancer – immune cell interactions and induction of tolerance mechanisms to develop actionable strategies that prevent immune evasion during the progression of breast cancer to lymph node metastasis.

Emerging evidence suggests that cancer cells utilize MHC-II induced tolerance mechanisms to evade anti-tumor immune responses, but the underlying mechanisms are not clear. A recent single cell analysis in pancreatic cancer identified a subset of cancer-associated fibroblasts expressing high levels of MHC-II without costimulatory molecules that may contribute to immune evasion of cancer cells^70^. In contrast to these observations, we recently demonstrated that epithelial stem cell MHC-II expression plays a critical role in promoting anti-tumor immune surveillance during intestinal tumor initiation^37^. Unlike breast cancer cells, intestinal stem cells exhibit expression of costimulatory-like molecules^37,38^. Furthermore, the differences in microenvironment of metLNs and intestine, including the microbiome, may account for the contrasting functional outcomes on anti-tumor immunity elicited by epithelial MHC-II expression. Future studies are needed to decipher the precise mechanisms through which MHC-II expression in different epithelial cancers influence tumorigenesis.

In summary, our study provides a comprehensive analysis of the progression of breast cancer to lymph node metastasis at single-cell resolution. These data help characterize the progression of the disease from the primary tumor to the lymph node and demonstrate how cancer cells avoid immune surveillance and survive in the lymph node—an organ in which priming of an anti-cancer immune response takes place^15,71^. Whether or not the cancer cell reprogramming of the lymph node microenvironment impacts any anti-cancer immune function, response to immunotherapy and outcomes for patients with lymph node metastasis requires further study.

## Acknowledgements

This work was financially supported by the grants from NIH (R21AI097745, R01CA214913, and R01HL128168 to T.P.P., P30CA045508-33 to S.B.), Rullo Family MGH Research Scholar Award (TPP), the Oliver S. and Jennie R. Donaldson Charitable Trust (S.B.), The Harold and Leila Y. Mathers Charitable Foundation (S.B), The Mark Foundation for Cancer Research (20-028-EDV to S.B.), Chan Zuckerberg Initiative / Silicon Valley Community Foundation (2021-239862 to S.B) and STARR Cancer Consortium (I13-0052 to S.B.). The authors would like to acknowledge Dr. Rakesh K. Jain, Dr. Yves Boucher and Dr. Michael Carroll for critical discussions. The authors would also like to thank CSHL Cancer Center Shared Resources (Single Cell Biology and Sequencing Technologies) supported in part by the NCI Cancer Center Support Grant 5P30CA045508 and Ulandt Kim from the NextGen core at Massachusetts General Hospital for the single-cell sequencing. We would also like to acknowledge Dr. Naxerova Kamila for providing the human breast cancer FFPE samples.

## Author Contribution

P.L., E.P., T.P.P. and S.B. conceived and designed the study and analyzed data. P.L., S.B., and T.P.P. wrote the manuscript, in which all coauthors commented on. E.P., D.J., P.L., P.A., J.W., S.C., Z.A., W.H., J.M.P, M.O., H.Z. and S.M. designed and performed experiments and analyzed data. S.C., Z.A., and W.H. performed and analyzed flow cytometry experiments. C.C., I.E., and S.B. helped with the single-cell sequencing experiment. P.A. and J.W. helped with the animal work. D.J. advised and helped with the manuscript. J.M.P helped with the breast cancer pathology data. A.S.K helped with human breast cancer pathology data quantification. P.H helped with animal experiment. All authors read and accepted the manuscript.

## Material and methods

### Mice and *in vivo* study

Six-to twelve-week-old, female Balb/c mice were housed in the E.L Steele Laboratories animal facility at Massachusetts General Hospital. 1×10^5^ 4T1 cells were injected into the fourth mammary fat pad (draining to the inguinal lymph node) in a volume of 50 μL per mouse. CYT387 (SelleckChem) was reconstituted in DMSO at 10mg/mL and diluted in injection buffer (5% PEG400, 5% Tween80 and 90% H_2_O) for oral gavage every two days of 25mg/kg for 7 doses. All animal experimental protocols were reviewed and approved by the Institutional Animal Care and Use Committee of the Massachusetts General Hospital, Boston, MA.

### UV-inactivated influenza virus PR8 inoculation

Influenza A virus (H1N1) A/Puerto Rico/8/34 (PR8) VR-95 and VR-95PQ were purchased from ATCC. Viruses were inactivated by UV light for 15 min at room temperature prior to injection. Mice were inoculated with 1×10^5^ PFU viruses + 2ug Anti-CD40 (BioXCell, Cat: BE0016-2) + 2ug poly I:C (InvivoGen, Cat: VAC-PIC) subcutaneously. On day 9-10, mice were euthanized and draining lymph nodes were collected.

### Cell lines

4T1, 4T1 sgRNA control cells, 4T1 *Ifngr1-/-, Ifngr2-/-, Ciita-/-*, B16F10, E0771, and MCa-P1362 cell lines were used in this study and tested to be mycoplasma free (MycoAlert-Lonza). Cells were cultured *in vitro* in Dulbecco’s modified Eagle’s medium (GIBCO, Invitrogen Life Technologies) supplemented with 10% (v/v) fetal bovine serum (Atlanta Biologicals). Cells were maintained in a 5% CO_2_-humidified incubator at 37^°^C.

### Immunohistochemical staining

Mouse inguinal lymph nodes were collected and embedded in OCT compound (Tissue-Tek) and frozen immediately on dry ice. For immunofluorescence staining, the lymph nodes were cut into 10 μm serial sections using a cryostat. Several tissue sections representing different depths of the lymph node were selected for staining. A hydrophobic pen was used to draw a circle around the tissues and then air-dried. Tissues were fixed by 4% formaldehyde at room temperature for 5 minutes. Slides were washed with 1xPBS twice and then blocked with 5% normal donkey serum + 0.5% TritonX-100 (Millipore Sigma) for 30 minutes at room temperature. Cytokeratin-FITC (Sigma, Cat: #F3418, 1:200 dilution) was used for the staining of all the cancer cells in metLNs. E-Cadherin (CST, Cat: #3195, Rabbit anti-mouse, 1:100 dilution) and vimentin (CST, Cat: #3932, Rabbit anti-mouse, 1:100 dilution) were used for staining of epithelial- and mesenchymal-cancer cells in the lymph node, respectively. These primary antibodies were diluted in IHC blocking buffer (1xPBS, 5% normal donkey serum, 0.5% TritonX-100) and incubated at 4^°^C overnight. After primary antibody staining, slides were washed with IHC washing buffer (1xPBS + 0.5% TritonX-100) 3-5 times for 5 minutes each. Anti-Rabbit-647 (1:200 dilution) was used for secondary antibody staining and incubated at room temperature for 1 hour. In all the slides, DAPI (1:1000) was used for nuclear staining at room temperature for 10-20 minutes. For cancer cells MHC-II staining, Cytokeratin-FITC (Sigma, Cat: #F3418, 1:200 dilution) was used for the staining of cancer cells in metLNs. Anti-mouse I-A/I-E-APC (Biolegend, Cat: #107614, clone M5/114.15.2, 1:100 dilution) was used for the MHC-II staining.

Formalin-fixed, paraffine-embedded deidentified human invasive ductal carcinoma specimens were obtained from the Massachusetts General Hospital Pathology department. For immunofluorescence and immunohistochemistry staining, the tissues were cut into a 5 μm thickness section and mounted on glass slides, followed by deparaffinization and heat-induced epitope retrieval in citrate-based antigen retrieval solution. MHC-II (LGII-612.14, CST, Catalog: 68258S) and Anti-human pan-Cytokeratin-Alexa647 (Biolegend, Catalog: 628604) were used for the staining. For MHC-II IHC staining, we used 1:800 dilution. For Cytokeratin staining, we used 1:100 dilution and incubated the primary antibody in cold room for overnight.

### Real-time PCR

Cells were administrated IFN-γ (10ng/ml, BioLegend, Cat: #575302) *in vitro* for 12h and 48h. mRNA was extracted by RNeasy plus kit (Qiagen) according to the manufacturer’s instructions. RNA reverse transcription was performed by iScript cDNA synthesis kits (Biorad). Real-time PCR mix with SYBR was purchased from Bio-Rad Laboratories. The primers for real-time PCR are listed below:

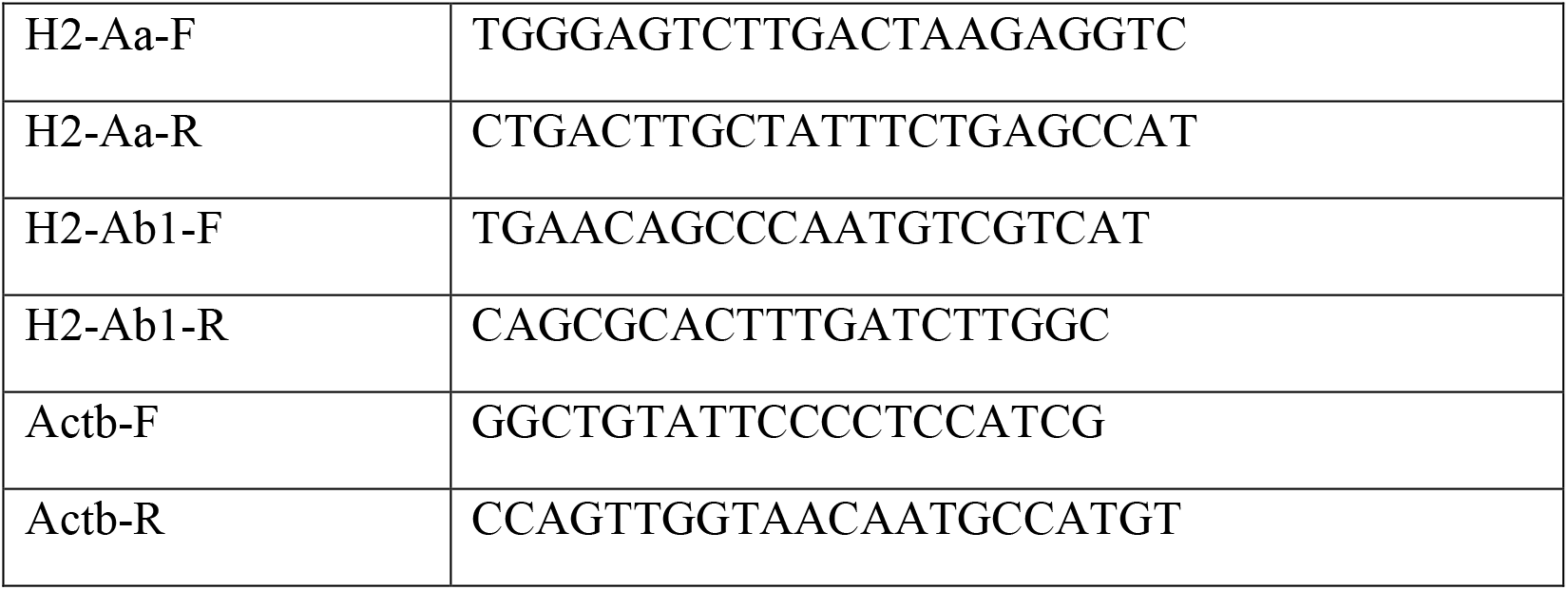

### CRISPR/Cas9-mediated gene disruption

sgRNA oligonucleotides for mouse *Ifngr1, Ifngr2* and *Ciita* genes were cloned into lentiCRISPRv2 according to the published protocol ^72^. For each gene disrupted, 1mg of the plasmid was transfected to 293T cells (ATCC #CRL-11268) using FuGENE 6 Transfection Reagent (Promega) to produce the retroviruses. The viral supernatants were harvested at 48 and 72 hours after transfection. To infect 1-2×10^5^ cells in a 6-well plate, we added 1 mL of viral supernatants with 2 mg/mL polybrene and added 1 mL fresh DMEM medium. Knockout clones were identified either by western blot or by flow cytometry analysis.

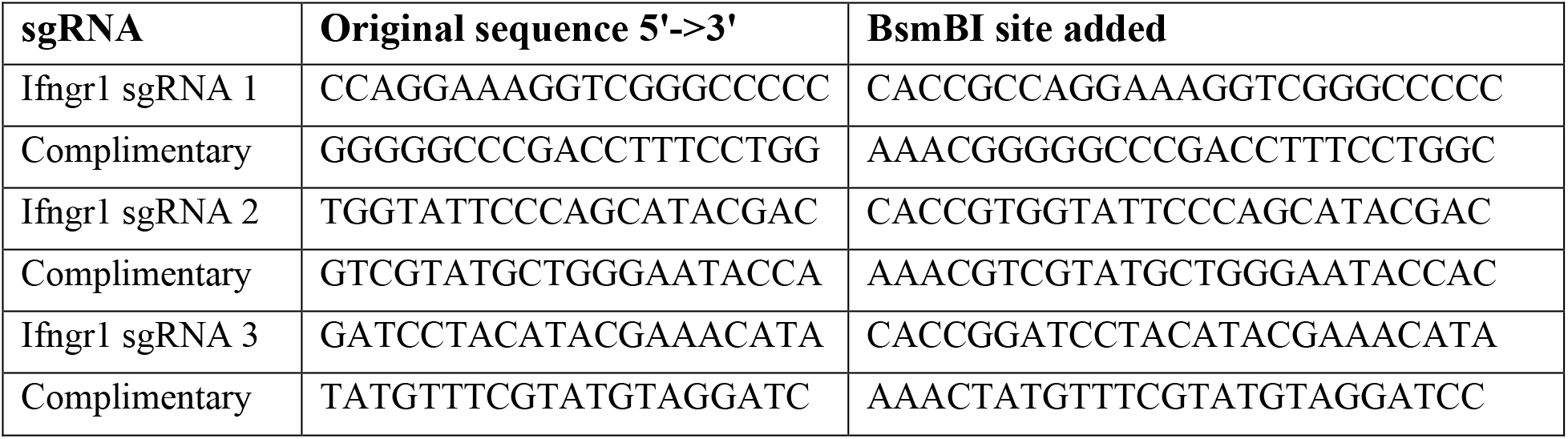
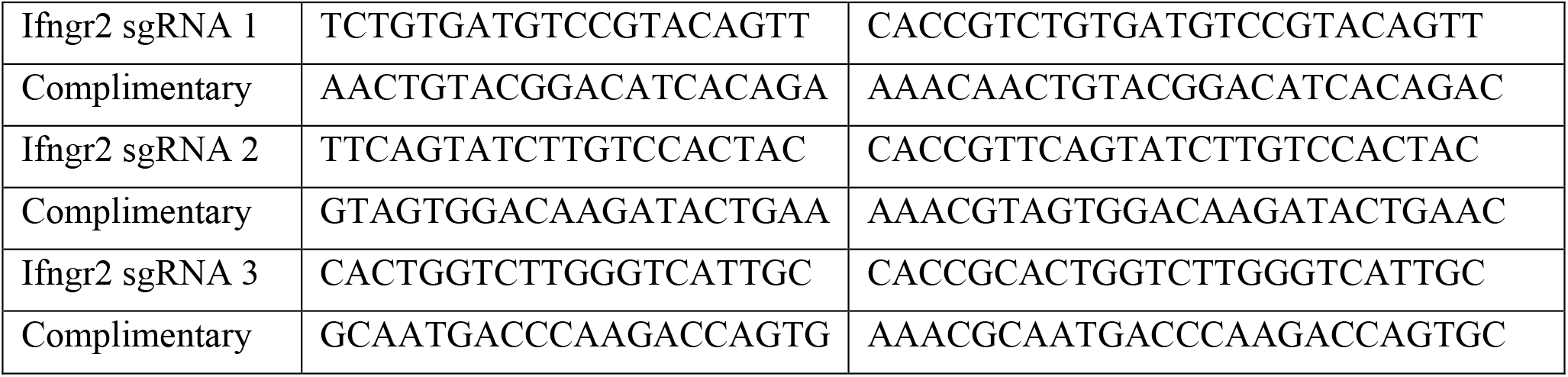

### Flow cytometry (cultured cells)

Cells were administrated IFN-γ (10ng/mL, BioLegend, Cat: #575302) *in vitro* for 24-48h. Cells were dissociated into single-cells by 0.05% Trypsin-EDTA (Gibco) at 37^°^C for 10 minutes. After dissociation, cells were filtered by 35μm strainer cap twice. Fc block was added to the pellet (1:500) and kept on ice for 15 minutes. After blocking, cells were stained with MHC-I (H-2Kd/Dd-Alexa-647, BioLegend, clone 34-1-25, 1:200 dilution) or MHC-II (I-A/I-E-APC-Cy7, BioLegend, clone M5/114.15.2, 1:200) in FACS buffer (PBS with 1% BSA) on ice for 20 minutes. After staining, cells were fixed, spun down and the cell pellet was resuspended in 150ul FACS buffer and analyzed by a BD LSR-II flow cytometer.

### Flow cytometry (mouse tissue)

We collected 4T1 metastatic inguinal lymph nodes from Balb/c mice. The lymph nodes were dissociated into single cells in digestion buffer (Collagenase P 0.2mg/mL, Diapase II 0.8mg/mL, DNase 0.1mg/mL, RPMI with 1% FBS) at 37^°^C. Every 5-8 minutes, tubes were agitated and then the contents were allowed to re-settle. Next, the supernatant was transferred to collecting buffer (RPMI with 1% FBS and 2mM EDTA). After 45-60 minutes, the lymph nodes were completely dissociated and centrifuged at 4 ^°^C, at 1200rpm for 5 minutes. After discarding the supernatant, Fc block was added to the pellet (1:500) and kept on ice for 15 minutes. After blocking, we stained with EpCAM-APC-eFluor780 (1:200 dilution) with FACS buffer (PBS with 1% BSA), leaving samples on ice in the dark for 20 minutes. After EpCAM staining, samples were washed by FACS buffer twice and then 1ml of fixation/permeabilization working solution was added to each sample at room temperature for 60 minutes. After fixation, 2mL of permeabilization buffer was added and the solution centrifuged at 1400 rpm for 5 minutes at room temperature. Antibodies for intracellular epitopes (cytokeratin-FITC, 1:100; vimentin-PE, 1:100) were then added and cells incubated at room temperature in the dark for 60 minutes. After intracellular staining, cells were washed 2-3 times by adding 2mL permeabilization buffer followed by centrifugation at 1400 rpm for 5 minutes at room temperature. The cell pellet was resuspended in 150 μL FACS buffer and analyzed by a BD LSR-II flow cytometer.

A separate cohort of mice was used to measure the presence of MHC-II molecules on cancer cells in 4T1 primary tumors and metLNs. Briefly, 14 days after tumor implantation we dissected the primary tumor and dissociated it into single cells as above. The mice were kept to further grow lymph node metastasis. At day 28, we sacrificed the mice and collected inguinal lymph nodes for flow cytometry. After dissociation, the red blood cells were lysed by ACK buffer (Thermo Fisher) if needed. Then, cells were filtered by 35 μm strainer cap twice. Fc block was added to the pellet (1:500) and kept on ice for 15 minutes. After blocking, cells were stained with MHC-II (I-A/I-E-BV605, BioLegend, 1:200), EpCAM-FITC (1:200) in FACS buffer (PBS with 1% BSA) on ice for 20 minutes. After cell surface marker staining, the cells were fixed and permeabilized for intracellular staining of vimentin-PE (1:100). After intracellular staining, cells were washed 2-3 times by adding 2mL permeabilization buffer followed by centrifugation at 1400rpm for 5 minutes at room temperature. The cell pellet was resuspended in 150ul FACS buffer and analyzed by a BD flow cytometer.

### Immune activation of lymph node

To measure T cell phenotype in metLNs, we set up four arms: saline, PR8, 4T1 and 4T1 + PR8. In the saline group, we subcutaneously injected 10 mL saline into the mice thigh of tumor naïve mice. In PR8 group, we inoculated 1×10^5^ PFU UV-inactivated PR8 influenza virus + 2μg Anti-CD40 (BioXCell, Cat: BE0016-2) + 2μg poly I:C (InvivoGen, Cat: VAC-PIC) via subcutaneous injection into the thigh of tumor naïve mice. 9-10 days later we collected the draining lymph nodes in both groups for flow cytometry. In the 4T1 and 4T1+PR8 groups, we implanted 4T1 cancer cells into 4^th^ mammary fat pad. When the tumor reached 250mm^3^, we dissected the primary tumor and randomized the mice into 2 groups. Five days later, we inoculated 1×10^5^ PFU UV-inactivated PR8 influenza virus + 2μg Anti-CD40 (BioXCell, Cat: BE0016-2) + 2μg poly I:C (InvivoGen, Cat: VAC-PIC) in half of the mice (4T1+PR8 group) and saline in the other half of the mice (4T1 group) via thigh injection. Another 9 days later, draining lymph nodes were collected for flow cytometry.

### Tissue collection and single-cell sequencing

4T1 cells were implanted into the left-side (supine position) fourth mammary fat pad of adult (6-10 weeks) female Balb/c mice. All analyzed lymph nodes were dissociated into single cells with digestion buffer (200 μg/mL collagenase I, 800 μg/mL Dispase, 100μg/mL DNase I). All samples used ACK buffer (Thermo Fisher) to lyse red blood cells and had the cell concentration and viability tested using a Nadia instrument (Dolomite Bio). The loading quantity of viable cells for 10X Genomics platform was estimated to capture around 5000 cells. In metLN1 and metLN2 dataset, we harvested the inguinal lymph nodes for single-cell sequencing from tumor bearing mice that had the primary tumor resected ∼ 2 weeks post implantation or when the tumors reached ∼500mm^3. We harvested the inguinal lymph nodes from naïve mice as control. Naïve LN1 and metLN1 were prepared and sequenced at the same time. Naive LN2 and metLN2 were prepared and sequenced at the same time. In the metLN2 data set, we further enriched the stromal cells by FACS cell sorting for CD45-population. Naïve LN1, Naïve LN2, metLN1 and metLN2 samples were sequenced with Illumina NextSeq High Output SE75 kit, with a format of 26×8×56. In the metLN3 dataset, we collected paired primary tumors and draining lymph nodes from the same mice (n = 4). The primary tumors were collected at day 14 or when tumor size reached ∼250mm^3^. After resection, we sutured the wound and allowed these mice to grow spontaneous lymph node metastases. The primary tumors were cut into 1-2 mm pieces and dissociated into single cells. To decrease bias from individual mice, we collected primary tumors from 4 mice and pooled them together for single-cell library preparation. To avoid sequencing reagent and sample loading bias, the primary tumor scRNA-Seq library was stored at -80^°^C until the metLN3 sample was available so that all sequencing occurred in the same flow-cell. On day 28, we sacrificed the mice and collected the inguinal lymph nodes (n = 4) to create the sample for the metLN3 dataset. In the metLN3 dataset, we enriched for tumor and stromal cells by MACS magnetic beads separation into CD45+ cells and CD45-cells with a CD45 Biotin antibody [10μL CD45-biotin antibody in 500uL MACS buffer (1xPBS, 1%FBS, 2mM EDTA)]. We estimated the proportion of cells from the lymph node and spiked-in 15,000 CD45+ cells into 1,000,000 CD45-cells. For 10X Genomics library preparation, we aimed to capture 5000 cells from the combined metLN3 sample. The metLN1 and metLN2 samples were sequenced independently. The paired primary tumor and metLN3 libraries were combined and loaded in the same flow-cell for sequencing on the Illumina HiSeq platform with High-Output mode PE26/98. Collectively, metLN1, metLN2 and metLN3 datasets after confirming the presence of cancer cells formed the data for the metLNs analyzed in this manuscript.

### Bioinformatic data analysis

#### UMAP and clustering

Cellranger (Version 3.0.2, https://support.10xgenomics.com/single-cell-gene-expression/software/overview/welcome) was used to pre-process the fastq-format raw data. Cellranger aggr was used to combine data from different runs. After that, Seurat^73,74^ (Version 3.1.3, https://satijalab.org/seurat/) was used to perform the graph-based clustering and analysis of differentially expressed genes. For quality control, we depleted cells with less than 200 genes detected as well as genes expressed in less than 3 cells. We also removed cells that had more than 10,000 featured RNAs and more than 20% mitochondrial genes. Cleaned data were normalized by NormalizeData function with the method LogNormalize. The most variable genes were detected by FindVariableFeatures function with the selection method “vst”. After normalization, data were scaled by ScaleData and RunPCA function was used to find the most significant principal components. For the primary tumor sample, we selected the top 50 PCAs for UMAP analysis and clustering. Using different clustering resolutions for the UMAP analysis, we found that c4 and c9 were deemed as one group at resolution 0.8 and 0.9, while c12 and c16 were grouped together at resolution 0.8. When the resolution increased from 1.0 to 1.3, we did not see additional cancer cell clusters. In contrast, additional myeloid populations were found when increasing resolution. We decided to use a resolution 1.0 for the downstream analysis as the cancer cell populations were well resolved. To normalize the batch bias of metLN samples, we employed Seurat integrated strategy ^74^. After integration, we selected the top 30 PCAs for UMAP analysis. The min.dist for UMAP analysis is 0.5 for both primary tumor and metLNs. For the cancer cell only analysis, we selected all cancer cell clusters in the primary tumor and metLNs, aggregated and integrated them. The top 20 PCAs were chosen for UMAP analysis and clustering.

#### Single-cell trajectory

Monocle ^28,29^ (version 2.10.1) was used to perform the trajectory analysis of cancer cells. Cell cycle regression of cancer cells was performed according to Seurat cell cycle scoring and regression method based on the canonical marker genes from Nestorowa *et al*. ^75^. The 2000 most variable genes were selected for the trajectory analysis.

#### GSVA analysis

For the GSVA analysis, msigdbr ^76^ (version 7.7.7) R package was used to retrieve the C2 curated gene sets from Molecular Signatures Database (MSigDB, https://www.gsea-msigdb.org/gsea/msigdb). GSVA ^77^ (version 1.30.0) R package was used for the single-cell gene set enrichment analysis. The limma ^78^ (version 3.38.3) package was used to identify the differential enriched gene sets across single cells clusters.

#### Human breast cancer single-cell sequencing analysis

The human breast cancer single-cell dataset GSE180286 was download from NCBI GEO database (https://www.ncbi.nlm.nih.gov/geo/). Seurat was used for the single-cell UMAP analysis. For quality control, we depleted cells with less than 200 genes detected as well as genes expressed in less than 3 cells. We also removed cells that had more than 10,000 featured RNAs and more than 20% mitochondrial genes. Sctransform (https://github.com/ChristophH/sctransform/) was used for data normalization and cell cycle regression. The cancer cell in each of these samples was identified by scGate (https://github.com/carmonalab/scGate) with gene signature (*KRT8+KRT8*+). We used UCell R package (https://github.com/carmonalab/UCell) to measure the single-cell gene signatures ^79^.

GO annotation and cell-cell interaction analysis: ShinyGO v0.61 (http://bioinformatics.sdstate.edu/go/) ^80^ was used for the gene ontology functional annotation, with p value cutoff 0.05 and top 30 most significant GO terms returned. CellChat ^44^ (version 1.4.0) R package was used for ligand-receptor analysis with the manually curated database of literature-supported ligand-receptor interactions in CellChatDB mouse.

### Imaging Processing

Stained lymph node sections were imaged by confocal microscopy (Olympus 1×81) using 10x air and 20x air objectives (Olympus). ImageJ (https://imagej.net/ImageJ) was used to process the multi-tiff format raw image data. Human breast cancer immunofluorescence images were analyzed in QuPath ^81^. For human breast cancer image data analysis, cells were identified based on a positive DAPI signal, and each of the cell populations were classified as positive or negative based on a single intensity threshold on mean expression within the cell. MHC-II+ cells located in both the tumor (Cytokeratin+) and stromal (Cytokeratin-) compartments were included in the quantitative analysis. The mean proportion of MHC-II+ Cytokeratin+ cells was subsequently calculated and reported.

### Statistical analysis

The unpaired t-test and one-way ANOVA was conducted across each group. An alpha value of 0.05 was considered statistically significant. All analyses were performed using Prism Version 9 Software (GraphPad).

## Data availability

Single-cell mRNA sequencing data generated to support this study have been deposited in NCBI GEO with GSE168181 accession number. The authors declare that all other data supporting the findings of this study are available within the paper and its supplementary information files.

## Antibodies and Reagents

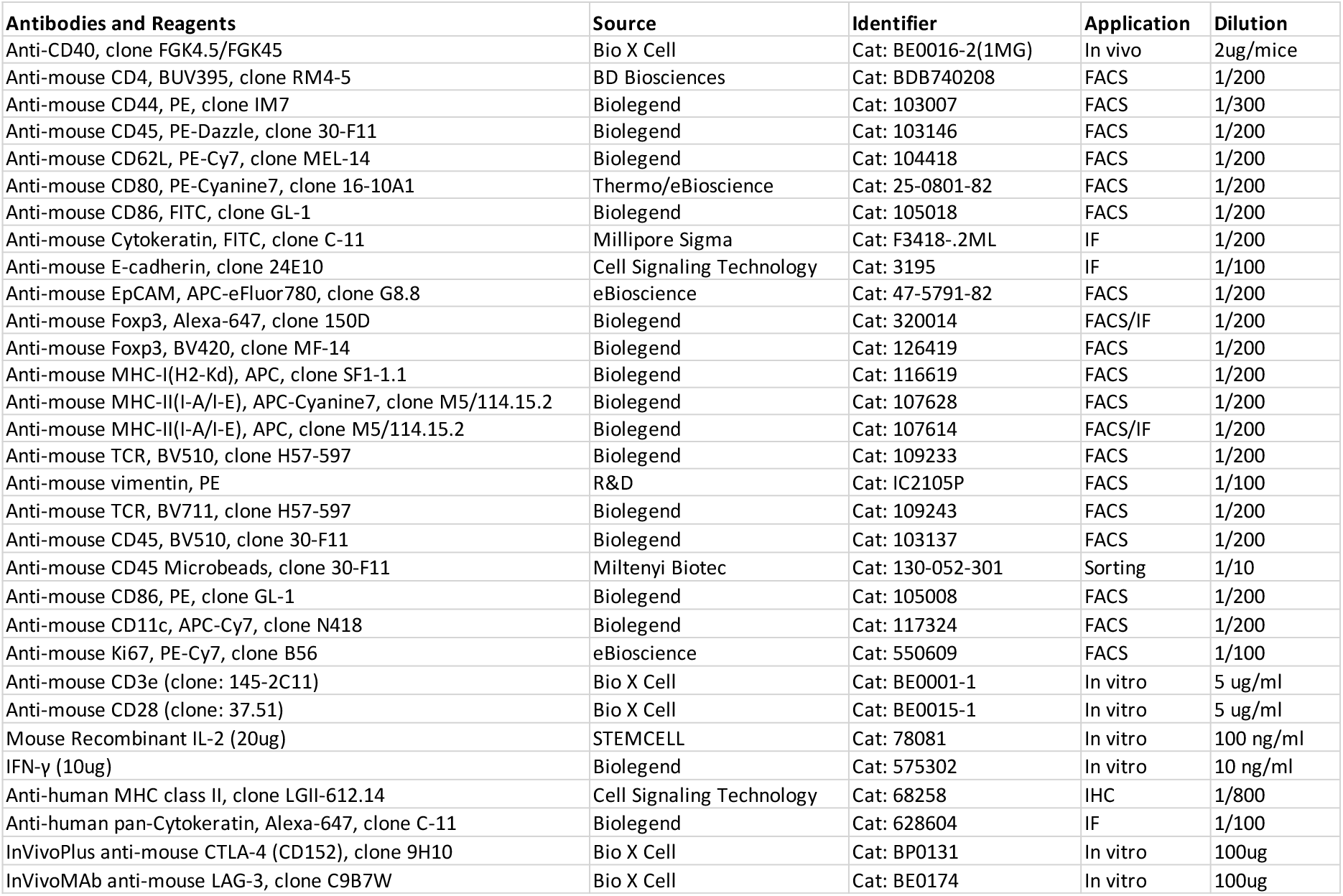

