## Supplemental figures for "Breast cancer progression and metastasis to lymph nodes reveals cancer cell plasticity and MHC class II-mediated immune regulation"

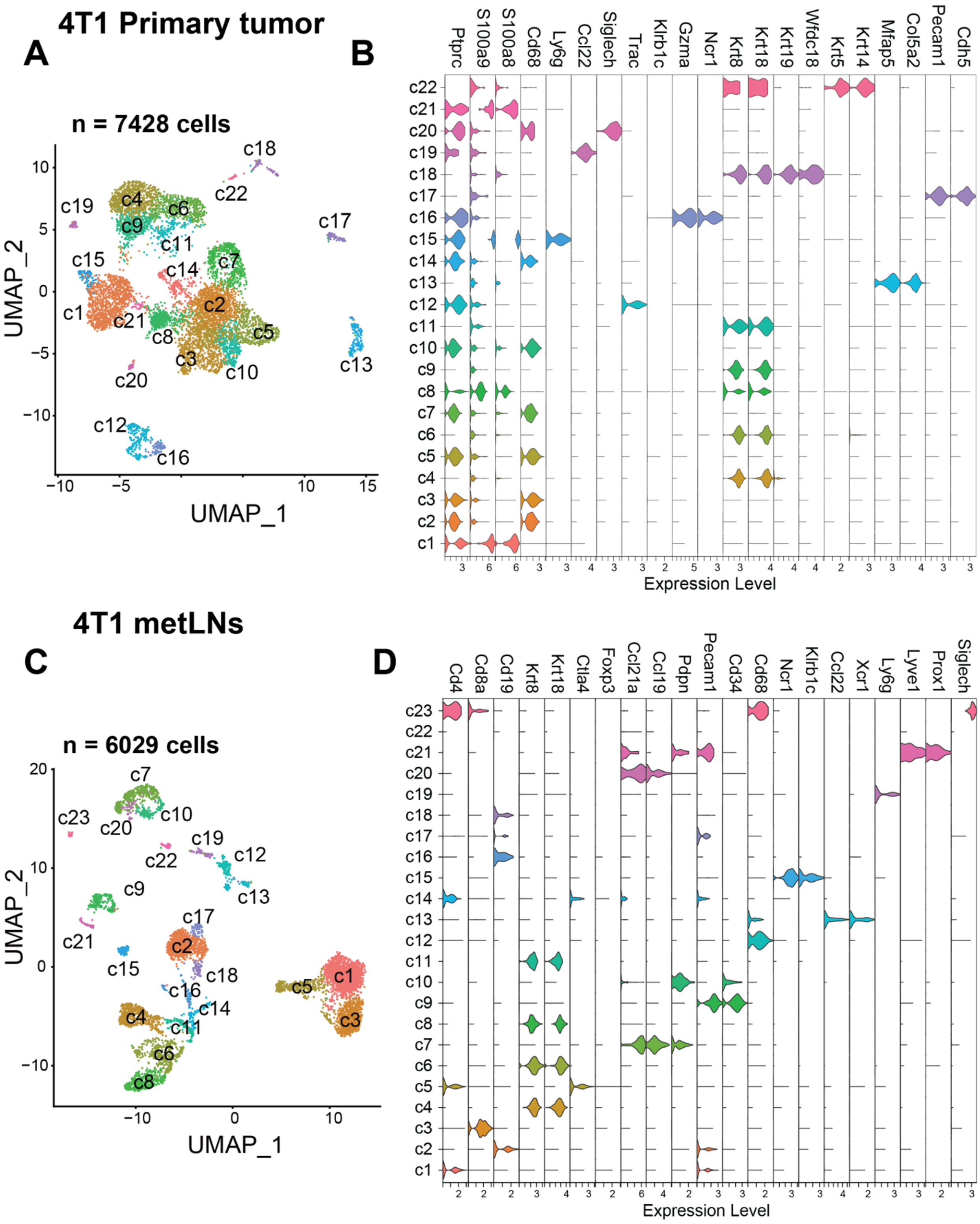
4T1 Primary tumor
A
n = 7428 cells
UMAP_2
UMAP_1
c1 c2 c3 c4 c5 c6 c7 c8 c9 c10 c11 c12 c13 c14 c15 c16 c17 c18 c19 c20 c21 c22
B
Ptprc S100a9 S100a8 Cd68 Ly6g Ccl22 Siglech Trac Klrb1c Gzma Ncr1 Krt8 Krt18 Krt19 Wfdc18 Krt5 Krt14 Mfap5 Col5a2 Pecam1 Cdh5
Expression Level
4T1 metLNs
C
n = 6029 cells
UMAP_2
UMAP_1
c1 c2 c3 c4 c5 c6 c7 c8 c9 c10 c11 c12 c13 c14 c15 c16 c17 c18 c19 c20 c21 c22 c23
D
Cd4 Cd8a Cd19 Krt8 Krt18 Ctla4 Foxp3 Ccl21a Ccl19 Pdpn Pecam1 Cd34 Cd68 Ncr1 Klrb1c Ccl22 Xcr1 Ly6g Lyve1 Prox1 Siglech
Expression Level

**Figure S1. The single-cell landscape of the 4T1 breast tumor and metLNs microenvironment.** **A)** The UMAP of 7428 cells from 4T1 primary tumor samples after correction for batch bias, colored by the clusters. We selected the top 50 principal components for the UMAP analysis with a minimum distance of 0.5 for the display. The resolution for the UMAP clustering is 1.0. **B)** Violin plots show the gene expression levels of selected marker genes in each cluster of cells from primary tumors. Gene expression values are log normalized. **C)** UMAP of 6029 cells from metLNs after correction for batch bias, colored by the clusters. metLN samples from three independent sequencing datasets were aggregated and normalized to correct for batch bias. We selected the top 30 significant principal components for UMAP analysis. The minimum distance for the UMAP is 0.5, the resolution for the UMAP is 1.0. **D**) Violin plots show the gene expression levels of selected marker genes in each cluster of cells from metLNs. Gene expression values are log normalized.

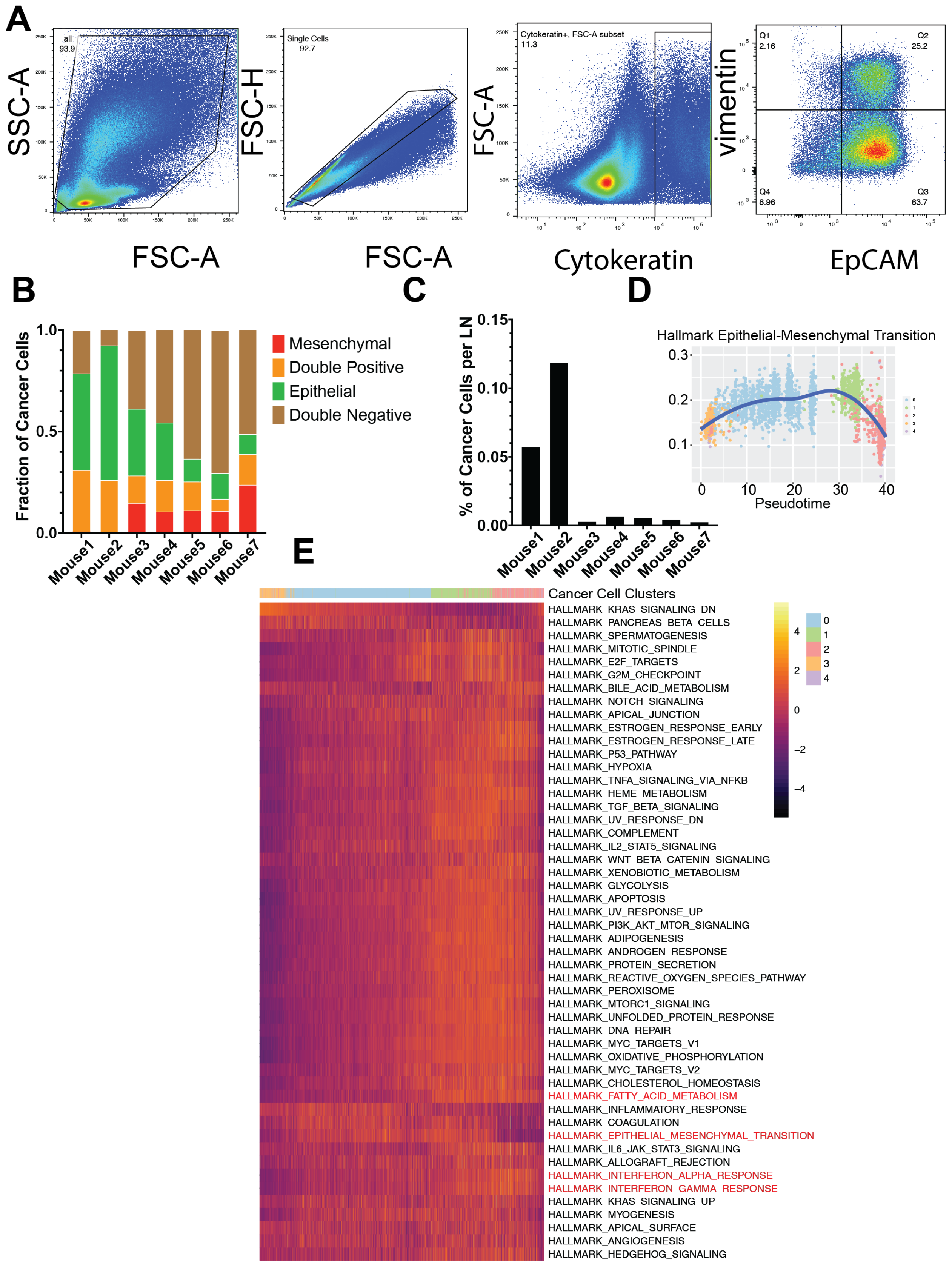
A
SSC-A
FSC-A
all
93.9
FSC-H
FSC-A
Single Cells
92.7
FSC-A
Cytokeratin
Cytokeratin+, FSC-A subset
11.3
vimentin
EpCAM
Q1
2.16
Q2
25.2
Q3
63.7
Q4
8.96
B
Fraction of Cancer Cells
1.0
0.5
0.0
Mouse1
Mouse2
Mouse3
Mouse4
Mouse5
Mouse6
Mouse7
Mesenchymal
Double Positive
Epithelial
Double Negative
C
% of Cancer Cells per LN
0.15
0.10
0.05
0.00
Mouse1
Mouse2
Mouse3
Mouse4
Mouse5
Mouse6
Mouse7
D
Hallmark Epithelial-Mesenchymal Transition
0.3
0.2
0.1
0
10
20
30
40
Pseudotime
E
Cancer Cell Clusters
HALLMARK_KRAS_SIGNALING_DN
HALLMARK_PANCREAS_BETA_CELLS
HALLMARK_SPERMATOGENESIS
HALLMARK_MITOTIC_SPINDLE
HALLMARK_E2F_TARGETS
HALLMARK_G2M_CHECKPOINT
HALLMARK_BILE_ACID_METABOLISM
HALLMARK_NOTCH_SIGNALING
HALLMARK_APICAL_JUNCTION
HALLMARK_ESTROGEN_RESPONSE_EARLY
HALLMARK_ESTROGEN_RESPONSE_LATE
HALLMARK_P53_PATHWAY
HALLMARK_HYPOXIA
HALLMARK_TNFA_SIGNALING_VIA_NFKB
HALLMARK_HEME_METABOLISM
HALLMARK_TGF_BETA_SIGNALING
HALLMARK_UV_RESPONSE_DN
HALLMARK_COMPLEMENT
HALLMARK_IL2_STAT5_SIGNALING
HALLMARK_WNT_BETA_CATENIN_SIGNALING
HALLMARK_XENOBIOTIC_METABOLISM
HALLMARK_GLYCOLYSIS
HALLMARK_APOPTOSIS
HALLMARK_UV_RESPONSE_UP
HALLMARK_PI3K_AKT_MTOR_SIGNALING
HALLMARK_ADIPOGENESIS
HALLMARK_ANDROGEN_RESPONSE
HALLMARK_PROTEIN_SECRETION
HALLMARK_REACTIVE_OXYGEN_SPECIES_PATHWAY
HALLMARK_PEROXISOME
HALLMARK_MTORC1_SIGNALING
HALLMARK_UNFOLDED_PROTEIN_RESPONSE
HALLMARK_DNA_REPAIR
HALLMARK_MYC_TARGETS_V1
HALLMARK_OXIDATIVE_PHOSPHORYLATION
HALLMARK_MYC_TARGETS_V2
HALLMARK_CHOLESTEROL_HOMEOSTASIS
HALLMARK_FATTY_ACID_METABOLISM
HALLMARK_INFLAMMATORY_RESPONSE
HALLMARK_COAGULATION
HALLMARK_EPITHELIAL_MESENCHYMAL_TRANSITION
HALLMARK_IL6_JAK_STAT3_SIGNALING
HALLMARK_ALLOGRAFT_REJECTION
HALLMARK_INTERFERON_ALPHA_RESPONSE
HALLMARK_INTERFERON_GAMMA_RESPONSE
HALLMARK_KRAS_SIGNALING_UP
HALLMARK_MYOGENESIS
HALLMARK_APICAL_SURFACE
HALLMARK_ANGIOGENESIS
HALLMARK_HEDGEHOG_SIGNALING
4
2
0
−2
−4
0
1
2
3
4

**Figure S2. The plasticity of cancer cells during lymph node metastasis.**
**A)** The representative flow cytometry gating strategy of EMT profiles of 4T1 cancer cells in metLNs. Cytokeratin (Alexa-488), EpCAM (APC-Cy7), vimentin (PE). **B)** Percentage of EpCAM+vimentin- (epithelial), EpCAM-vimentin+ (mesenchymal), EpCAM-vimentin- (double negative), and EpCAM+vimentin+ (double positive) cancer cells compared to all cancer cells (cytokeratin+) in metLNs. **C)** Percentage of cancer cells in metLNs measured by FACS. **D**) The single-cell enrichment score of MSigDB Hallmark Epithelial-Mesenchymal Transition gene set. Cancer cells were ranked by pseudotime, and the blue line represents the loess regression of the enrichment score. **E)** The single-cell gene sets enrichment analysis of cancer cells in 4T1 primary tumor and metLNs. The gene sets are from GSEA MSigDB hallmark gene sets. Each column represents a single cell, each row represents a gene set. Cancer cells were ranked by the pseudotime. Yellow represents high, purple represents low.

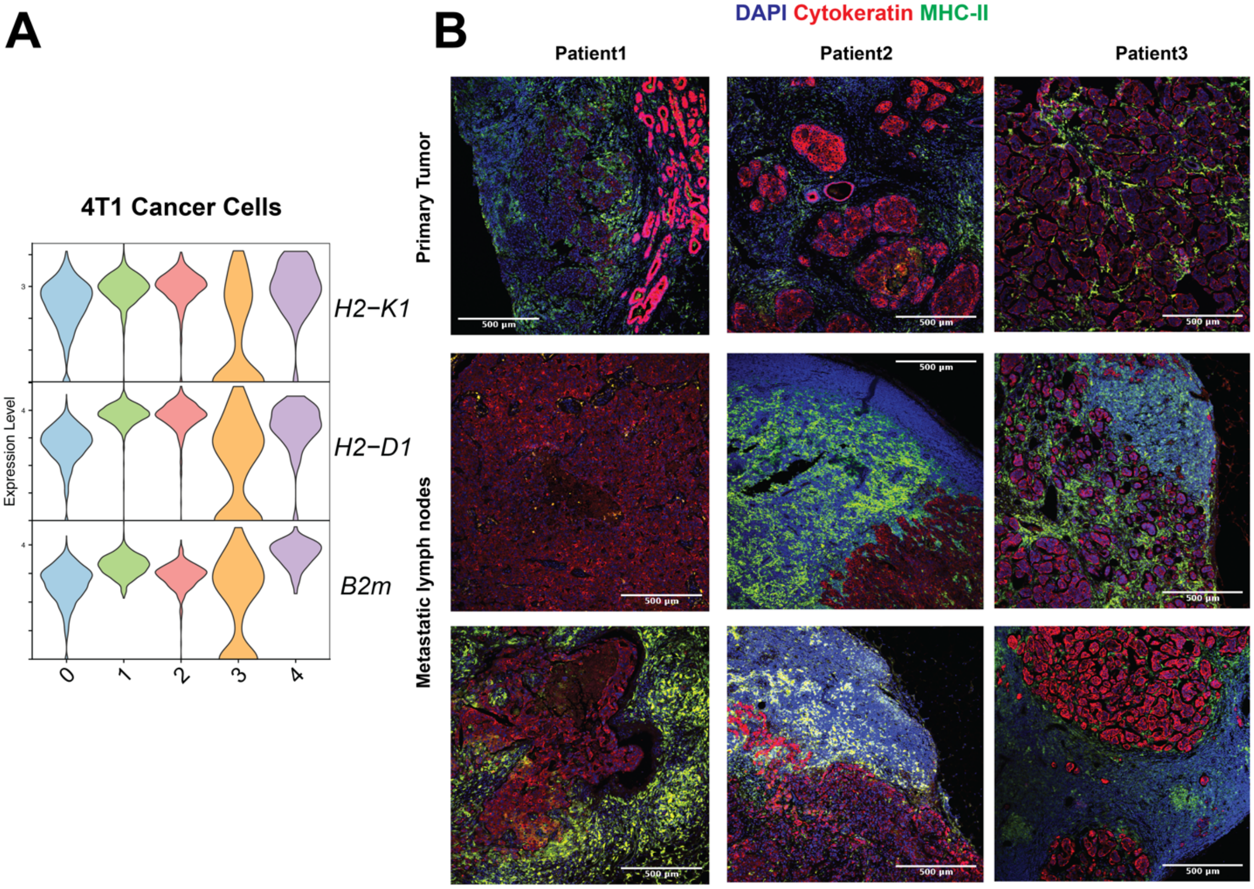


**Figure S3. The presence of MHC-I in 4T1 cancer cells and human breast tumors MHC-II staining.**

**A**) The violin plot shows the gene expression of MHC-I molecules in 4T1 cancer cells (clusters 1, 2, 4 from metLNs). **B**) Immunofluorescence staining of nuclei (DAPI, blue), pan-cytokeratin (red) and MHC-II (green) in human breast tumors and metastatic lymph nodes in FFPE specimens. The thickness of the sections is 5 µm. The deidentified breast tumor samples were provided by Massachusetts General Hospital pathology department.

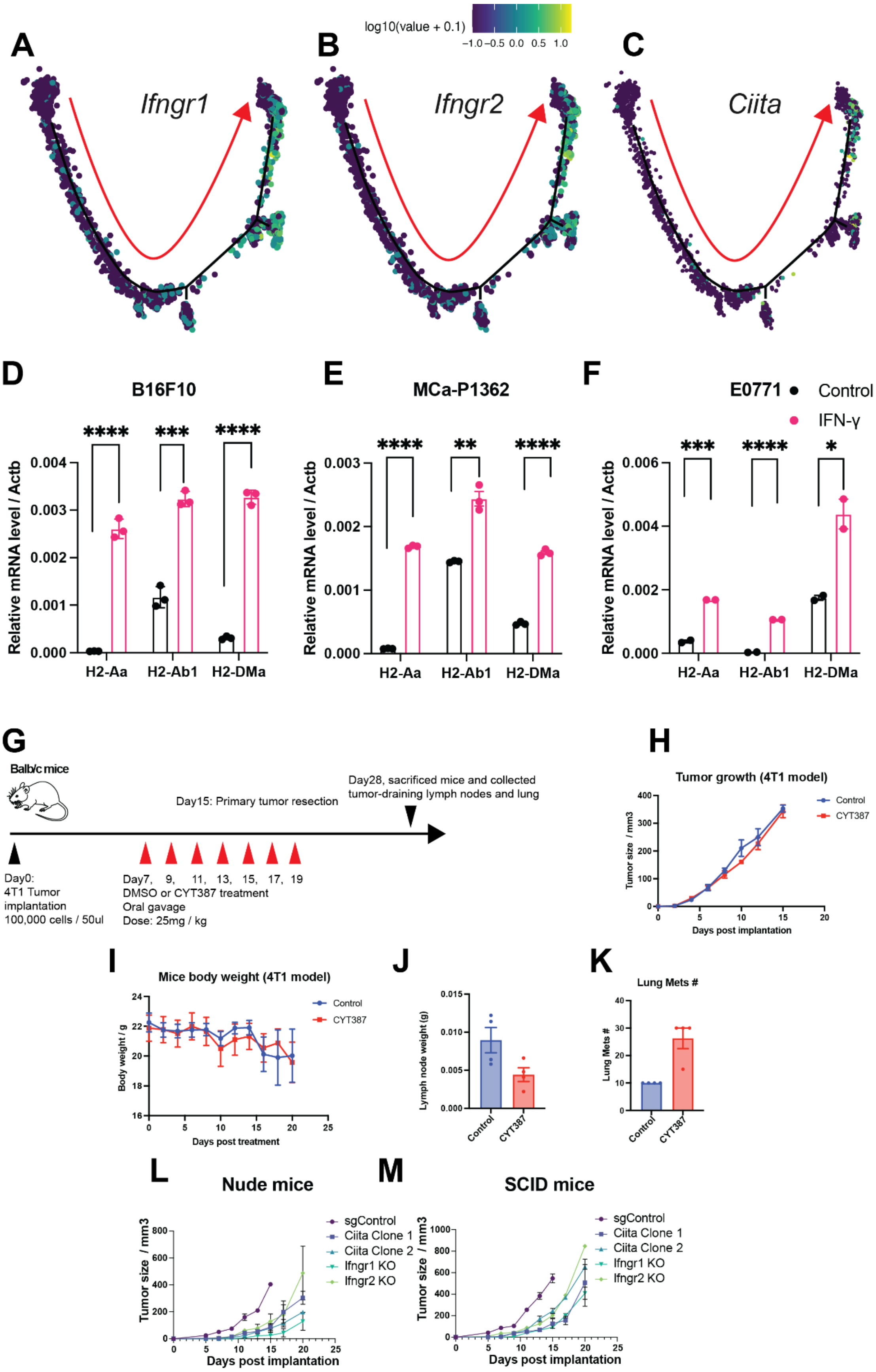
A
B
C
log10(value + 0.1)
−1.0 −0.5 0.0 0.5 1.0
Ifngr1
Ifngr2
Ciita
D
E
F
B16F10
MCa-P1362
E0771
Control
IFN-γ
Relative mRNA level / Actb
H2-Aa
H2-Ab1
H2-DMa
****
***
**
*
G
Balb/c mice
Day0:
4T1 Tumor
implantation
100,000 cells / 50ul
Day7, 9, 11, 13, 15, 17, 19
DMSO or CYT387 treatment
Oral gavage
Dose: 25mg / kg
Day15: Primary tumor resection
Day28, sacrificed mice and collected tumor-draining lymph nodes and lung
H
Tumor growth (4T1 model)
Tumor size / mm3
Days post implantation
Control
CYT387
I
Mice body weight (4T1 model)
Body weight / g
Days post treatment
J
Lymph node weight (g)
K
Lung Mets #
L
Nude mice
M
SCID mice
sgControl
Ciita Clone 1
Ciita Clone 2
Ifngr1 KO
Ifngr2 KO
Tumor size / mm3
Days post implantation

**Figure S4. Targeting Interferon-gamma pathway controls tumor growth.**
**A-C)** The gene-expression of *Ifngr1*, *Ifngr2*, and *Ciita* were projected to the single-cell trajectories. Gene expression values are scaled, and log normalized. The red arrow indicates the pseudotime trajectory of cancer cells progression. **D-F**) IFN-γ induced expression of MHC-II molecules *H2-Aa*, *H2-Ab1* and *H2-DMa in vitro*. B16F10 (melanoma), MCa-P1362 (breast cancer) and E077 (breast cancer) cells were treated with or without IFN-γ (10 ng/mL) for 24 hours. Student's t-test was used for the statistical analysis. *, $p$-value $< 0.05$; **, $p$-value $< 0.01$; ***, $p$-value $< 0.001$; ****, $p$-value $< 0.0001$. **G)** The experiment design of JAK/STAT inhibitor CYT387 *in vivo* assay. **H)** 4T1 tumor growth in Control and CYT387 treated group (n=4). **I)** Mice body weight change in Control and CYT387 treated group (n=4). **J)** The weight of the tumor-draining lymph nodes at day 28. **K)** The number of pulmonary metastases at day 28. **L-M)** The tumor growth of 4T1 sgRNA control cells, *Ifngr*1/2 and *Ciita* knockout cells in immunodeficient nude mice (n=4) (L) and SCID mice (n=4) (M).

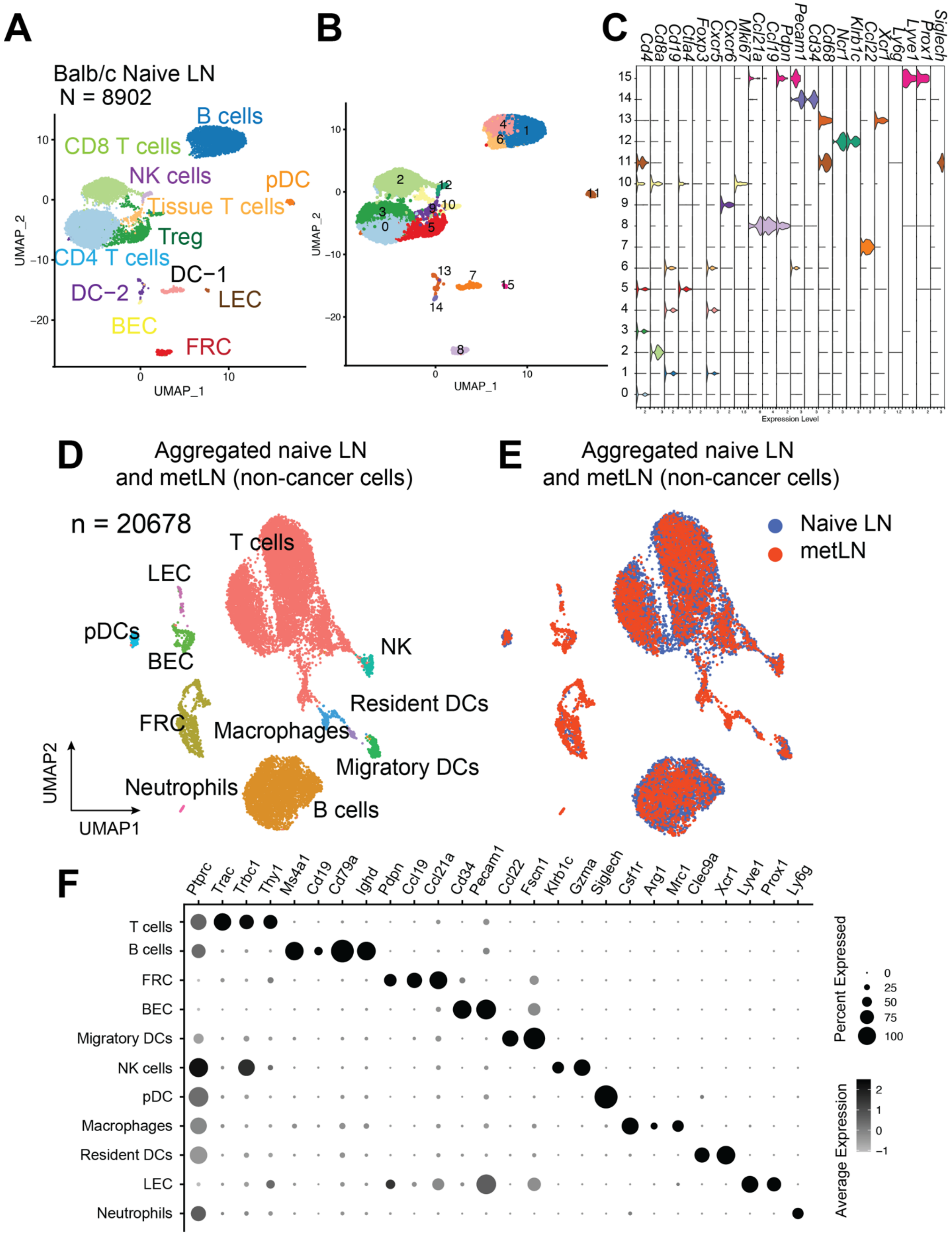
A
Balb/c Naive LN
N = 8902
B cells
CD8 T cells
NK cells
pDC
Tissue T cells
Treg
CD4 T cells
DC−1
DC−2
LEC
BEC
FRC
UMAP_1
UMAP_2
B
C
Cd4
Cd8a
Cd19
Ctla4
Foxp3
Cxcr5
Cxcr6
Mki67
Ccl21a
Ccl19
Pdpn
Pecam1
Cd34
Cd68
Ncr1
Klrb1c
Ccl22
Xcr1
Ly6g
Lyve1
Prox1
Siglech
Expression Level
D
Aggregated naive LN and metLN (non-cancer cells)
n = 20678
T cells
LEC
pDCs
BEC
NK
FRC
Resident DCs
Macrophages
Migratory DCs
Neutrophils
B cells
UMAP1
UMAP2
E
Aggregated naive LN and metLN (non-cancer cells)
Naive LN
metLN
F
Ptprc
Trac
Trbc1
Thy1
Ms4a1
Cd19
Cd79a
Ighd
Pdpn
Ccl19
Ccl21a
Cd34
Pecam1
Ccl22
Fscn1
Klrb1c
Gzma
Siglech
Csf1r
Arg1
Mrc1
Clec9a
Xcr1
Lyve1
Prox1
Ly6g
T cells
B cells
FRC
BEC
Migratory DCs
NK cells
pDC
Macrophages
Resident DCs
LEC
Neutrophils
Percent Expressed
Average Expression

**Figure S5. The single-cell atlas of inguinal lymph nodes in naïve Balb/c mice and metLNs.** **A-B)** The UMAP of 8902 cells from inguinal lymph nodes in tumor-free Balb/c mice (naïve mice), (A) colored by the cell types, (B) colored by the clusters. We selected the top 30 principal components for the UMAP analysis with a minimum distance of 0.5 for the display. The resolution for the UMAP clustering is 0.8. **C)** Violin plots show the gene expression levels of selected marker genes in each cluster of cells from naïve lymph node. Gene expression values are log normalized. **(D)** The UMAP of aggregated cells in naïve lymph nodes and 4T1 metLNs (n = 20678). Top 30 principal components were chosen for the UMAP analysis, with minimum distance of 0.5 and clustering resolution of 0.8, and cells were colored by main cell types in the lymph nodes. **(E)** The UMAP of aggregated cells in naïve lymph nodes and 4T1 metLNs grouped by the samples. **(F)** The gene-expression pattern of selected marker genes in the main cell types in the lymph nodes. Black represents high expression; gray represents low expression. The size of the circle represents the proportion of cells expressing the indicated genes in each cluster.

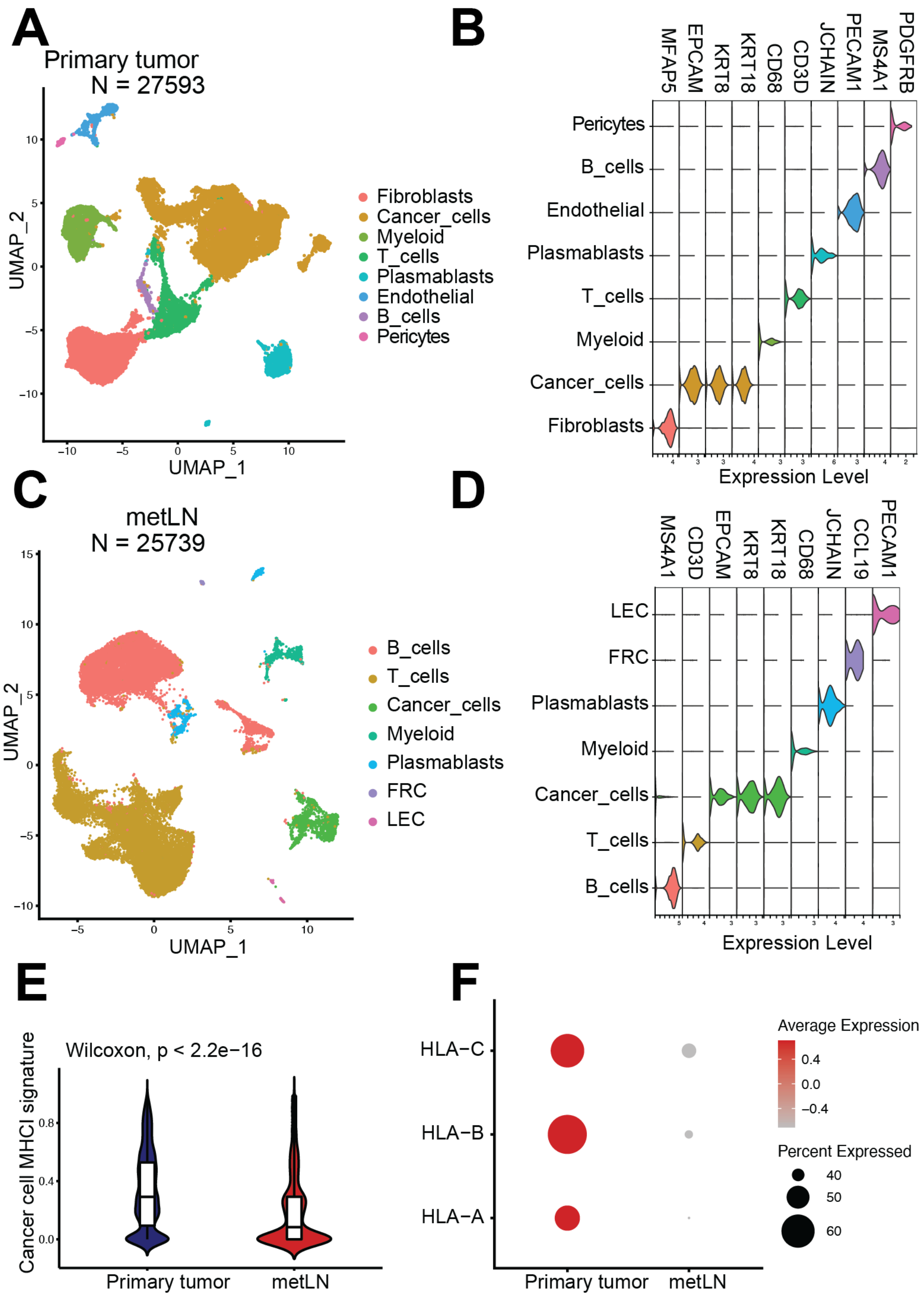
A
Primary tumor
N = 27593
UMAP_2
UMAP_1
Fibroblasts
Cancer_cells
Myeloid
T_cells
Plasmablasts
Endothelial
B_cells
Pericytes
B
MFAP5
EPCAM
KRT8
KRT18
CD68
CD3D
JCHAIN
PECAM1
MS4A1
PDGFRB
Pericytes
B_cells
Endothelial
Plasmablasts
T_cells
Myeloid
Cancer_cells
Fibroblasts
Expression Level
C
metLN
N = 25739
UMAP_2
UMAP_1
B_cells
T_cells
Cancer_cells
Myeloid
Plasmablasts
FRC
LEC
D
MS4A1
CD3D
EPCAM
KRT8
KRT18
CD68
JCHAIN
CCL19
PECAM1
LEC
FRC
Plasmablasts
Myeloid
Cancer_cells
T_cells
B_cells
Expression Level
E
Wilcoxon, p < 2.2e−16
Cancer cell MHCI signature
Primary tumor
metLN
F
HLA−C
HLA−B
HLA−A
Primary tumor
metLN
Average Expression
0.4
0.0
−0.4
Percent Expressed
40
50
60

**Figure S6**. **The single-cell atlas of human breast cancer and metLNs.**
**A**) The UMAP of 27593 cells from human breast tumors from NCBI GEO database GSE180286. We selected the top 30 principal components for the UMAP analysis with a minimum distance of 0.5 for the display. The resolution for the clustering is 0.5. **B**) Violin plots show the gene expression levels of cell types-specific marker genes in breast tumors. **C**) The UMAP of 25739 cells from breast cancer metastatic lymph nodes from NCBI GEO database GSE180286. We selected the top 30 principal components for the UMAP analysis with a minimum distance of 0.5 for the display. The resolution for the clustering is 0.5. (**D**) Violin plots show the gene expression levels of cell types-specific marker genes in metLNs. **E**) Violin plots show the MHC-I gene signature in cancer cells. **F**) The expression profiling of MHC-I genes in cancer cells.
